# Understanding structural and functional diversity of ATP-PPases using protein domains and functional families in CATH database

**DOI:** 10.1101/2023.10.12.562014

**Authors:** Vaishali P. Waman, Jialin Yin, Neeladri Sen, Mohd Firdaus-Raih, Su Datt Lam, Christine Orengo

**Author notes:** contributed equally (Vaishali P. Waman, Jialin Yin).

## Abstract

ATP-Pyrophosphatases (ATP-PPases) are the most primordial lineage of the large and diverse HUP (<u>H</u>IGH-motif proteins, <u>U</u>niversal Stress Proteins, ATP-<u>P</u>yrophosphatase) superfamily. There are four different ATP-PPase substrate-specificity groups, and members of each group show considerable sequence variation across the domains of life despite sharing the same catalytic function. Over the past decade, there has been a >20-fold expansion in the number of ATP-PPase domain structures most recently from advances in protein structure prediction (e.g. Alphafold2). Using the enriched structural information, we have characterised the two most populated ATP-PPase substrate-specificity groups, the NAD-synthases (NAD) and GMP synthases (GMPS). We performed local structural and sequence comparisons between the NADS and GMPS from different domains of life and identified taxonomic-group specific structural functional motifs. As GMPS and NADS are potential drug targets of pathogenic microorganisms including *Mycobacterium tuberculosis*, structural motifs specific to bacterial GMPS and NADS provide new insights that may aid antibacterial-drug design.

## Introduction

ATP-**P**yrophosphatase (ATP-PPase), together with **H**IGH-motif Proteins and **U**niversal Stress Protein, constitute the HUP domain superfamily (CATH ID:3.40.50.620)^1^. They have been proposed to share a common ancestry, supported by their shared topology. The HUP core consists of a 3-layer αβα sandwich, with a 5-stranded parallel β-sheet flanked by 2 α-helices on each side, resembling a Rossman-fold domain^1,2^. In addition, ATP-PPase and HIGH-proteins use nucleotide-based molecules as their cofactors or co-substrates, similar to many Rossman-fold proteins^3^. These nucleotide-binding domains retain a Phosphate-Binding Loop (PBL) for co-substrates like ATP, and are characterised by an SGGxDS motif^4^.

ATP-PPases catalyse a two-step ligation process. In the first step, ATP is hydrolysed into AMP and a pyrophosphate ion, and the AMP binds to a Substrate A, forming an adenylated intermediate. In the second step, a nitrogen-group-containing Substrate B binds to the intermediate, which then undergoes nucleophilic attack^5–9^ (**Equation.1**) This results in the formation of product A-B, and the release of AMP^2^. Phylogenetic studies of HUP domains^1^ and comparisons of the reaction schemes across the HUP superfamily, have suggested that ATP-PPase are the primordial lineage of HUPs^2^. Therefore, studying the diversity within ATP-PPase lineage may provide a more detailed insight into the evolutionary history of HUP domain structures.

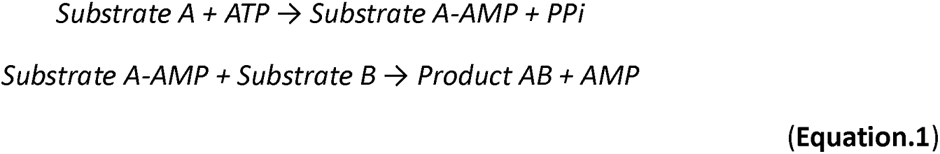

A previous study of the HUP superfamily revealed the presence of multiple substrate-specificity groups within the ATP-PPase lineage^10^. Enzymes within each specificity group recruit their unique substrate A, thereby synthesizing various products^10,11^. These groups include the Asparagine Synthase (ASNS)^12^, Argininosuccinate Synthase (ASS)^9^, Guanosine Monophosphate Synthase (GMPS)^6^ and Nicotinamide Adenine Dinucleotide Synthase (NADS)^13^. Interestingly, despite variations in substrate A, ASNS, GMPS, and NADS all utilize ammonia as their substrate B (**Table.1**).

**Table.1.**
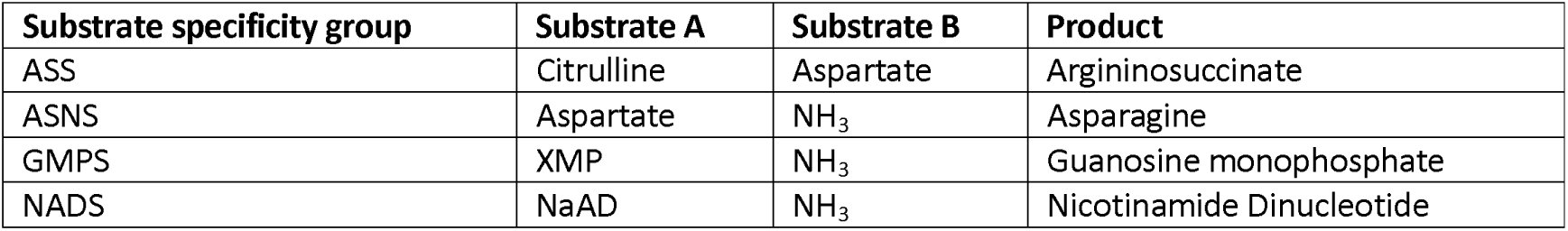
The four substrate specificity groups of the ATP-PPase lineage in the HUP superfamily.

Furthermore, all ASNS, all GMPS and some NADS are referred to as Glutamine-dependent bifunctional enzymes with two catalytic domains^14^. In addition to the common ATP PPase domain, they possess a unique counterpart domain which generates NH_3_ as the second substrate of the ATP-PPase domain reaction. These counterpart domains hydrolyse a Glutamine into a Glutamate and an NH_3_ molecule^14^. ASNS exploit their N-terminal domain^12^, GMPS have a Glutamine Amidotransferase domain^6^ (GATase) and Glutamine-dependent NADS (Gln-NADS) have a Carbon Nitrogen hydrolase domain^15^ (CN). Associated with their different Multidomain Architectures (MDA), their mechanisms of transporting NH_3_ from their Gln-hydrolysing site to the ATP PPase-domain catalytic site also varies. ASNS and Gln dependent NADS have both been observed to have continuously present NH_3_-tunnels^12,15^, whereas GMPS has been proposed to transiently form an NH_3_-tunnel *via* an intrinsic disordered loop called the Lid region^6,7,16^. Some exceptional NADS recruit an ammonium ion from the cell environment and are single-domain enzymes^13^. While ATP-PPase domains of these enzymes share the same global structures, we explored the evolution of certain regions of the ATP-PPase domains that accommodate different NH_3_/NH_4_^+^ binding and transporting mechanisms.

This is now possible since there are sufficient ATP-PPase structures and sequences for characterizing their functional regions. Over the past decade, the number of experimentally determined and classified structures within the HUP superfamily increased by more than 20-fold. In 2010, the CATH domain structure classification v3.2 contained only 85 HUP domain structures, with 12 of them identified as ATP-PPases^10^. Notably, these ATP-PPases were exclusively derived from prokaryotic origins. In the most recent release of the CATH database (version 4.3)^17^, the number of experimentally determined ATP-PPase structures has risen to 74, out of a total of 1,685 HUP domains. Furthermore, the revolutionary AlphaFold2 algorithm^18,19^ has enabled the prediction of structures for a vast number of HUP superfamily sequences, encompassing 41,313 sequences. High-quality AlphaFold2 predicted models (>90 pLDDT score) can be predicted for a non-redundant subset (at 90% sequence identity) of 3,637 HUP sequences^20^. Collectively, these structures, derived from diverse species across all domains of life, give a robust foundation for our investigation.

The HUP superfamily in CATH v4.3 has been further classified into 922 Functional Families (FunFams) through sequence comparisons^21,22^. Each FunFam comprises a set of sequence relatives whose residue conservation patterns differ from those present in other FunFams^23^, suggesting different substrate specificity or oligomerisation. Of these, four FunFams are associated with GMPS functionality, and 11 ATP-PPase domain structures have been experimentally determined. Although members of these FunFams perform similar catalytic functions, they originate from distinct taxonomic groups. Similarly, there are six FunFams associated with NADS and two FunFams associated with ASNS. We explored these different substrate-specificity groups to determine whether different FunFams within them evolved distinct modes of domain synergism^24^ with their counterpart Gln-hydrolysing domains, whilst possessing similar catalytic functions^5,25,26^.

Drawing on the enriched structural information and a range of bioinformatic tools, we performed detailed classification and characterization of the GMPS and NADS FunFams. Structural comparisons between four GMPS FunFams showed that the Lid region is differentially conserved between FunFams, supported by previous experiments^7,27,28^. I n addition, sequence comparisons between GMPS FunFams exposed several residues within the Lid region that are differentially conserved, providing new insights into various modes by which the Lid region regulates Gln-hydrolysis and NH_3_-transportation. Similarly, for the Gln-dependent NADS, our computational analysis highlighted differentially conserved residues and potential allosteric sites that may affect FunFam-specific NH_3_-transportation mechanisms that have not yet been experimentally investigated. Besides functional determination, these comparisons allowed us to make inferences on the evolutionary history of the Gln-dependency of the two most populated ATP-PPase substrate-specificity groups in the HUP superfamily.

By analysing the predicted functional regions and differentially conserved residues, we identified structural motifs and conserved residues, specific to each FunFam. Knowledge of structural motifs specific to bacterial GMPS/NADS FunFams could facilitate anti-bacterial drug development, as GMPS and NADS are essential enzymes involved in de novo purine synthesis^29^ and the production of metabolic coenzyme NAD^30^, respectively. These enzymes represent potential targets for drug development against pathogenic microorganisms, such as *Mycobacterium tuberculosis* and Staphylococcus aureus^15,31–36^. The ASSAM (Amino acid pattern Search for Substructures and Motifs) algorithm^37^ allowed us to search these structural motifs against several protein structure databases including CATH and AlphaFold2, thus verifying the specificity and high conservation of the bacterial-FunFam-specific structural motifs.

## Results and Discussion

### ATP-PPases comprise 15% of the HUP Superfamily

Structural and sequence data of ATP-PPase domains were gathered from the HUP superfamily (CATH: 3.40.50.620) of the CATH database V4.3^23^. The HUP superfamily encompasses a total of 41,313 sequences, which have been further classified into 922 FunFams^21^. There were 39 FunFams identified as ATP-PPases based on their conserved Phosphate Binding Loop motif SGGxDS^2^ and their Gene Ontology (GO)^38^ and Enzyme Commission (EC) annotations. They account for 6,168 sequences in total, which is about 15% of the HUP superfamily (**Supplementary.Table1**).

Among the 39 ATP-PPase FunFams, 15 of them have experimentally determined structures available in the Protein Data Bank (PDB)^39^. For the sequences belonging to an additional 3 FunFams with no experimental structures but high sequence diversity^40^ (DOPS >70) that allowed for the detection of evolutionarily conserved residues, we predicted structures using AlphaFold2 (AF2)^18,19^. A multiple structural alignment of the 75 structures from the 18 FunFams with structural data, was constructed by mTM-align^41,42^. The structural alignment guided UPGMA^43^ phylogenetic tree clearly supports the presence of four substrate-specificity groups within ATP-PPases (**Table.2; Supplementary.Figure1**)^10^.

**Table.2.** The ATP-PPase subgroups have different substrate specificities.

| Substrate-specificity Group | Step-one Substrate Specificity | Step-two substrate source | EC Number |
| --- | --- | --- | --- |
| Asparagine synthase (ASNS) | Aspartate | Ntn-domain (CATH: 3.60.20.10) | 6.3.5.4 |
| Argininosuccinate synthase (ASS) | Citrulline | Cell environment | 6.3.4.5 |
| GMP synthase<br>(GMPS) | Xanthosine monophosphate<br>(XMP) | GATase domain<br>(CATH: 3.40.50.880) | 6.3.5.2 |
| NAD synthase<br>(NADS) | deamido-NAD<sup>+</sup> (NaAD) | Cell environment<br>or<br>CN domain<br>(CATH: 3.60.110.10) | 6.3.1.5<br>or<br>6.3.5.1 |

### A loop adjacent to the catalytic site distinguishes GMPS and NADS

We examined the Glutamine-dependent ATP-PPases, the GMPS and NADS, which have sufficient structural information for detailed structural analyses. By applying the Zebra3D^44^ program to the 60 representative structures from the four FunFams annotated as GMPS and the six NADS FunFams, we identified a Specificity-Determining Region (SDR) (**Figure.2a**) that exhibits different structural conservation between GMPS and NADS. This SDR corresponds to the region between the 4^th^ and 5^th^ β-strands of the common core structure within the HUP superfamily. In NADS, the SDR is a fully conserved loop involved in substrate Nicotinic acid adenine dinucleotide (NaAD) binding^13,44,45^. In GMPS, the SDR is the intrinsically disordered Lid region^6,7,28^ that does not directly interact with the substrate. Conformational change of this region is thought to facilitate the functioning of GMPS in several ways^7,16,46^, discussed further below.

**Figure.2.**
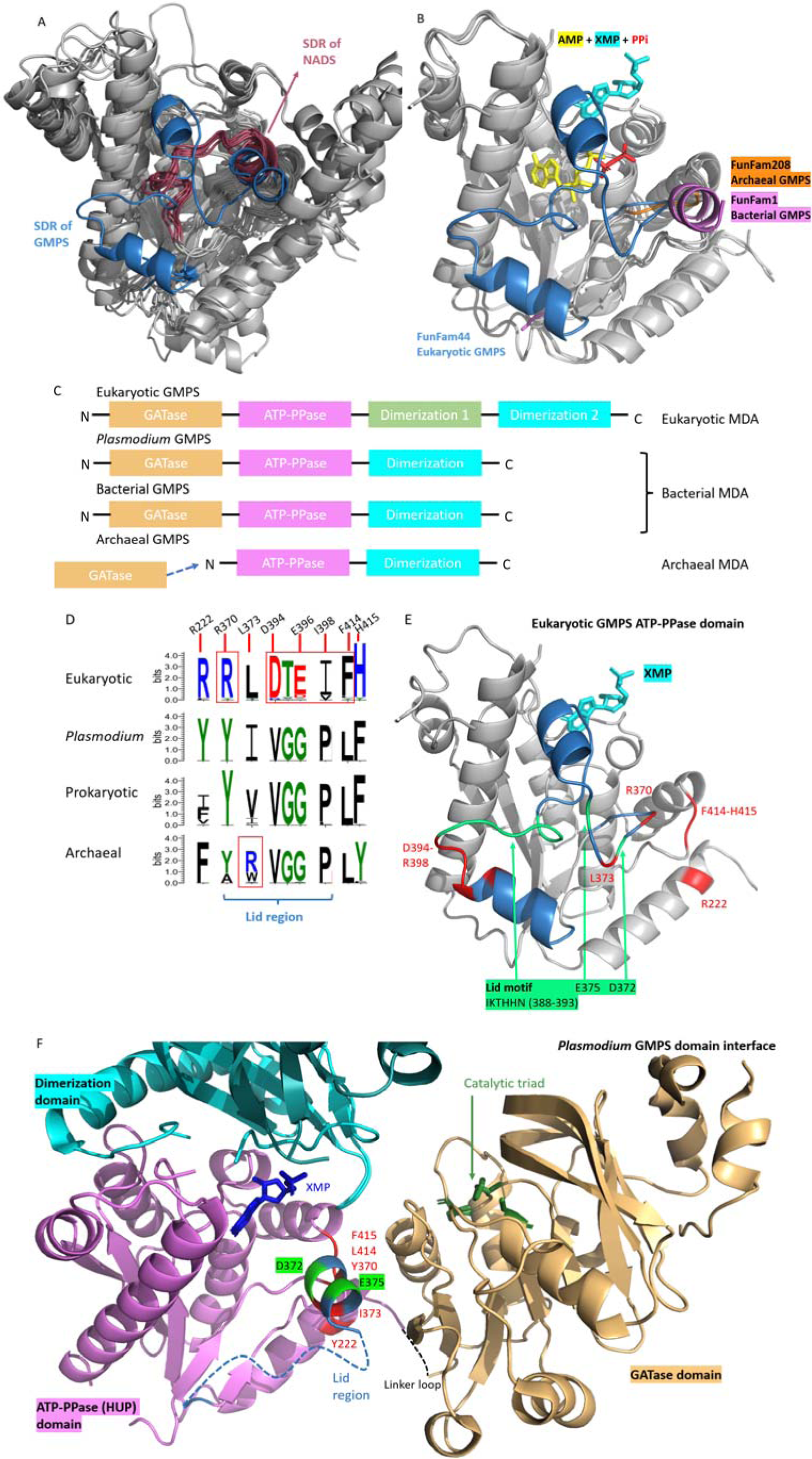
Features that distinguish different FunFams of GMPS. (a) Zebra3D identified regions that distinguish the GMPS (blue) and NADS (dark red). (b) Zebra3D subclassification of GMPS based on the Lid loop structures. (c) GroupSim identified Specificity Determining Positions (SDPs) and their varying residue conservation patterns within GMPS FunFams. The size of the residue symbols indicates their level of conservation, the colours represent their biochemical properties. The figure was generated using WebLogo3^51,52^. (e) SDP residues labelled on the eukaryotic GMPS ATP-PPase domain crystal structure (PDB: 2vxo). Red: SDP residues. Green: residues conserved by all GMPS. (f) SDP and fully conserved residues labelled on Plasmodium GMPS. The GATase domain catalytic site (dark green) is known from the literature^6^.

Although the Lid region is only fully resolved in eukaryotic GMPS, Zebra3D was able to classify the GMPS ATP-PPase domains into taxonomic subgroups based on the partially resolved sections of the prokaryotic Lid region (**Figure.2b**). This subclassification is similar to that of the FunFams, which distinguishes different domains of life (**Table.2**).

**Table.2.**
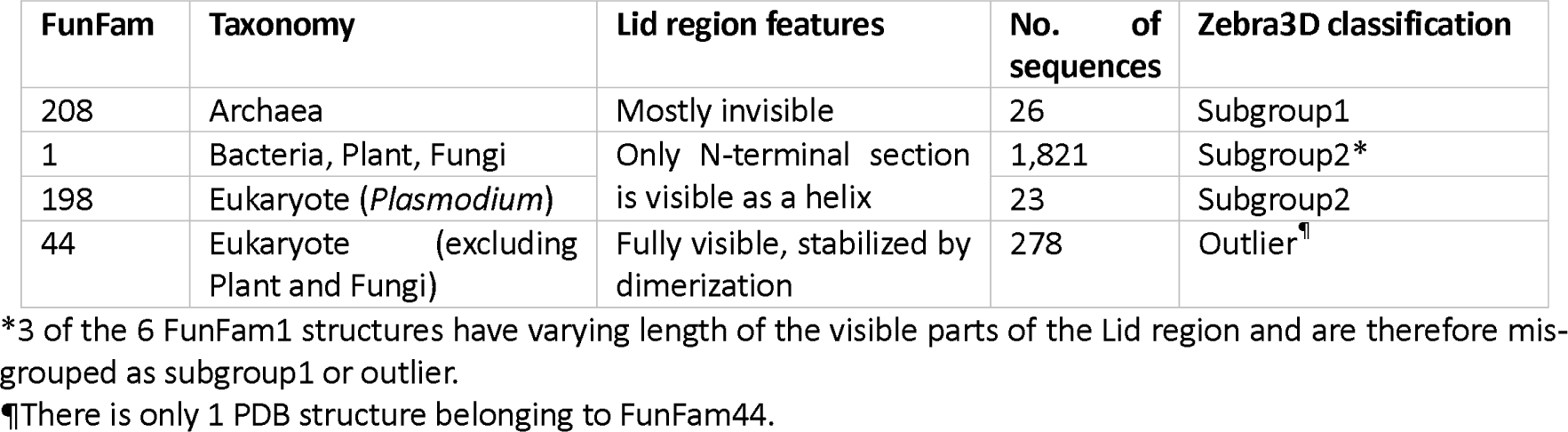
Zebra3D structural subclassification of the GMPS is comparable to that of the sequence based GMPS FunFam subclassification.

### Structural variation in the GMPS Lid-region structural is associated with different Multidomain Architectures

GMPS enzymes typically consist of three domains: the N-terminal Glutamine Amidotransferase domain (GATase), the ATP-PPase domain, and a C-terminal dimerization domain (**Figure.2c**). The exceptions are all from the archaeal GMPS FunFam (FunFam208), where the ATP-PPase and GATase subunits exist as separate polypeptides. These subunits transiently interact with each other to form a holoenzyme^27,47^. As a result, the Lid region of the apo-form archaeal ATP-PPase subunits (residues 132-153 on the representative structure, PDB:3a4i) is exposed on the surface of the ATP-PPase domain, resulting in low resolution and Zebra3D subclassification as a separate subgroup (**Figure.2b, Table.2**).

For the members of FunFam1 and the FunFam198, the Lid region is at the domain interface between the ATP-PPase and GATase domains. FunFam1 is dominated by bacterial GMPS (97.4%) and FunFam198 contains entirely unicellular parasitic GMPS from genus *Plasmodium*. The N-terminal sections of their Lid regions are stabilized by their interaction with GATase domains^48^. Zebra3D analysis was able to detect this feature of the *Plasmodium* GMPS and classified them into subgroup2 (**Table.2, Figure 2b**).

FunFam44 is dominated by eukaryotic GMPS with only one experimentally determined structure available (PDB: 2vxo). It exhibits a distinct Multidomain Architecture (MDA) compared to prokaryotic GMPS enzymes, owing to the presence of an additional dimerization domain (**Figure.2a**). In this case, the eukaryotic Lid region is located on the domain interface between the ATP-PPase and the additional dimerization domain of another protomer of the homodimer^28^. This unique interaction results in the fully stabilized Lid region structure (**Figure.2b**).

The variation in the flexibility of the Lid region, as evidenced by their absence/presence in the crystal structure, is linked to the different MDAs of GMPS across different FunFams, suggesting that GMPS enzymes from different domains of life may employ specific types of ATP-PPase-GATase domain synergism^49^, probably *via* different conformational changes of their Lid region^16^ (discussed below).

### Sequence analysis of GMPS ATP-PPases supports the regulatory and catalytic roles of the Lid region

GroupSim sequence analysis identified Specificity Determining Positions (SDPs) specific to the four GMPS FunFams and corroborating the Zebra3D subclassification. At these positions, each FunFam tends to have unique residue-conservation patterns^50^. This sequence analysis overcomes the analytical challenge posed by the flexible prokaryotic Lid regions, whose structures were only partially resolved by crystallography. To avoid ambiguity, the GMPS residues are all numbered according to the only fully resolved structure, the human GMPS (PDB: 2vxo).

For the four FunFams of the GMPS, a total of nine SDPs were identified, six of which are within the Lid region (**Table.3; Figure 2d**). Two of the remaining three are spatially close to the Lid region (within 5Å distance) with the remaining SDP close to these two (**Figure.2e**). The functional roles of three of these 9 SDP residues in *Plasmodium falciparum* G M P S (PfGMPS, PDB: 3uow) have been investigated previously^7^. Y222, Y370, and F415 of PfGMPS are situated at the interface between the ATP-PPase domain and the GATase domain (**Figure.2f**). These hydrophobic and aromatic SDP residues were found to engage in hydrophobic interactions with residues in the GATase domain^7^. This suggests that the flexible Lid region may interact with and regulate the GATase domain by changes in its conformation^7,16^ (details in the following section).

**Table.3.** GroupSim identified Specificity Determining Positions across the 4 GMPS FunFams.

| Residue (2vxo) | GroupSim score | Eukaryotic | Plasmodium | Prokaryotic | Archaeal | Feature |
| --- | --- | --- | --- | --- | --- | --- |
| 222 | 0.74 | R | Y* | I/F/V | F | Domain-domain interface* |
| 373 | 0.74 | L | I | V | R/W | Between catalytic residues D372, E375 |
| 415 | 0.75 | H | F* | F | Y | Domain-domain interface*, 4Å to lid region |
| 414 | 0.86 | F | L | L | L | 3.5Å to lid region |
| 370 | 0.91 | R | Y* | Y | Y/A | Domain-domain interface* |
| 394 | 0.96 | D | V | V | V | One residue away from Lid motif |
| 395 | 0.93 | T | G | G | G | Two residues away from Lid motif |
| 396 | 0.86 | E | G | G | G | Three residues away from Lid motif |
| 398 | 0.83 | I | P | P | P | Five residues away from Lid motif |

Red: SDPs with different conservation patterns across the four GMPS FunFams.
Purple: SDPs where only eukaryotic GMPS have a unique conservation pattern.
\*Annotations confirmed from a study of *Plasmodium falciparum* GMPS<sup>7</sup>.

In addition to the SDPs, some catalytic residues conserved in all GMPS FunFams are found within the Lid region. These residues are D372, E375, and a six-residue motif (residues 388-393) known as the Lid motif (IK(T/S)HHN)^28^ (**Figure.2e, f**), which is detected by the Scorecons algorithm^40^. The residues D372, E375, K389, H391 and H392^7,28,48^ are involved in the first step of GMP production (**Equation.1**), which is adenylating the substrate XMP to generate XMP-AMP^2,49^. However, these catalytic residues are distant from the XMP-binding site in all the resolved GMPS structures^7,27,28,46,47^. This suggests a potential common function of the Lid region in delivering the catalytic residues D372, E375 and the Lid motif to the ATP-PPase domain catalytic site^7,16,46,48^.

### Different Multidomain Architectures reflect different modes by which the Lid region regulates ATP-PPase-GATase domain synergism

Although the Lid region structures and the Multidomain Architectures (MDA) vary across the GMPS FunFams, activity assays suggest that all GMPS-ATP-PPase domains/subunits are able to activate the GATase domain Gln-hydrolysis reaction by catalysing XMP-adenylation^49,53,54^. It is possible that the Lid region first delivers the catalytic residues to the ATP-PPase active site for XMP-adenylation^46^, and then undergoes further conformational change, relocating to free up space in the active site for GMP production. As suggested by previous studies^7,46,48^, this movement away from the active site may allow the Lid region to interact with the GATase domain active site, and subsequently activate Gln-hydrolysis. While the Lid region function in XMP-adenylation could be common for all GMPS, the subsequent GATase domain activation may vary amongst the domains of life, as suggested by their specific MDAs and SDPs (**Figure.3**).

**Figure.3.**
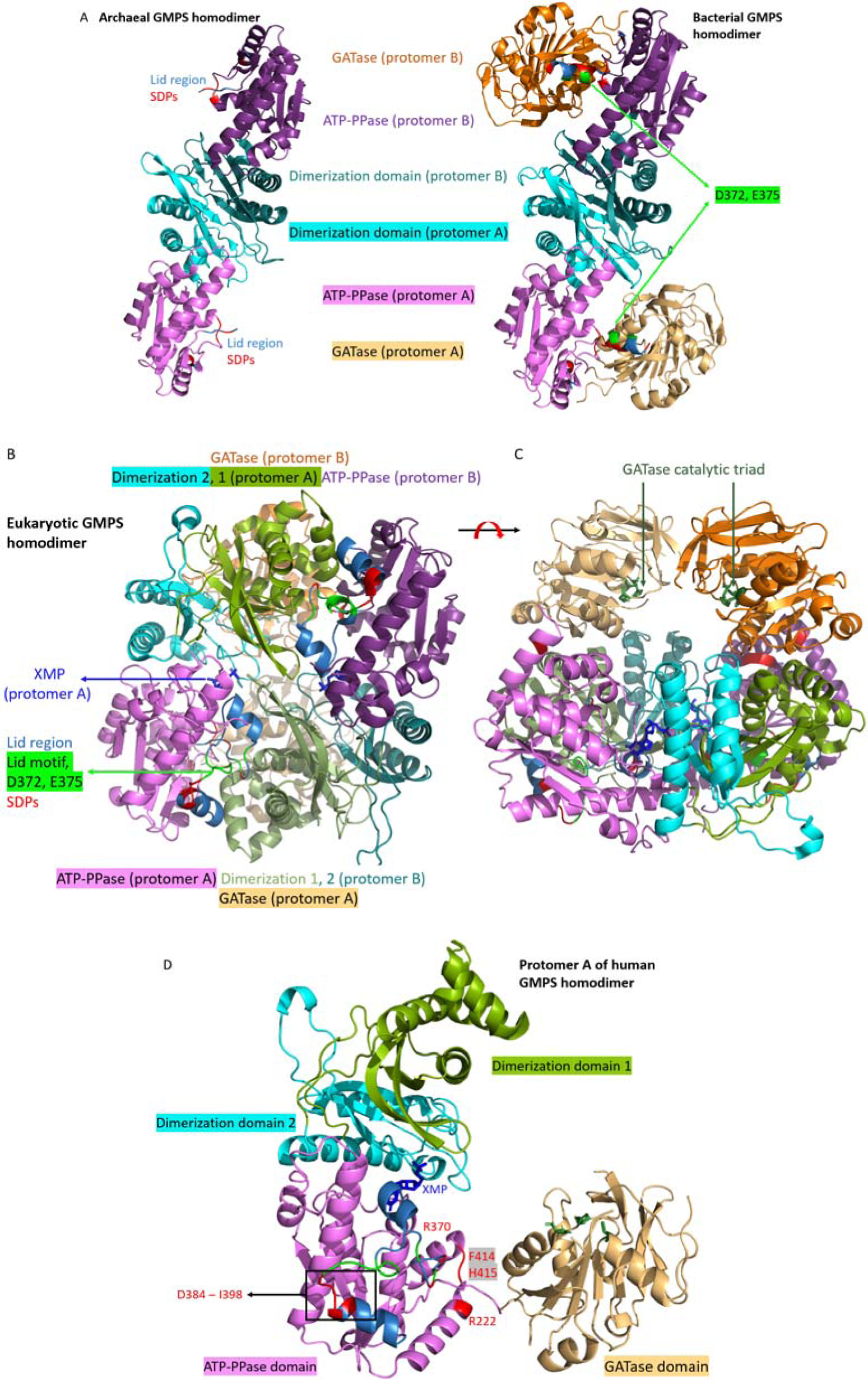
Different Multidomain Architectures corresponding to different modes of GMPS domain synergism. (a) Left: Archaeal dimer of ATP-PPase subunits (PDB: 3a4 i); Right: bacterial and Plasmodium have similar GMPS dimers. (b) Bottom and side views of the eukaryotic GMPS dimer (PDB: 2vxo). (c) Protomer A of eukaryotic GMPS. Colour scheme: Light blue – Lid region; Red – SDP residues; Green – conserved catalytic residues and motifs.

Movement of the Lid region to facilitate XMP adenylation may also contribute to the rotation of the GATase domain^7^. Molecular dynamics studies^7^ have shown that the GATase domain of *Plasmodium* GMPS rotates and positions its catalytic triad close to the ATP-PPase catalytic site; this is confirmed by a crystallographic structure of a GATase-domain-rotated *Plasmodium* GMPS (PDB: 4 wio)^7^. When only AMP is bound (*E.coli* GMPS, PDB: 1g pm)^6^, or XMP bound (*Plasmodium* GMPS, 3uow)^46^ their structures are similar to the apoenzyme. Therefore it is likely that both XMP and ATP need to be present at the ATP-PPase domain catalytic site to initiate the GATase domain rotation^7^. *Plasmodium* GMPS SDP residues Y222 and Y370 are involved in hydrophobic interactions with the GATase domain on the unrotated PfGMPS (**Figure.2f**). After the rotation, Y370 and F415 are involved in the ATP-PPase-GATase hydrophobic interface instead^7^. Participation of Y370 in this domain-domain interaction emphasizes the critical role of the Lid region in GMPS domain synergism.

Bacterial FunFam members not only share SDP residue conservation patterns with the *Plasmodium* GMPS (**Table.3**), but also possess the same MDA as the *Plasmodium* FunFam (**Figure.2c**). They function as homodimers with a Z-shape planar arrangement, where the ATP-PPase domains are on the elbows and the GATase domains are on the two ends (**Figure.3a**)^6^. Furthermore, the specific domain arrangement supports the Lid region in its cascade of movements to catalyse XMP-adenylation first and then facilitates the activation of GATase domain for Gln-hydrolysis and rotation of the GATase domain to support the subsequent transportation of NH_3_ back to the ATP-PPase domain for the last step of GMP production. Since *Plasmodium* are simple unicellular eukaryotes and the Bacterial FunFam is more populated and diverse than the *Plasmodium* one, we propose that this MDA and mode of Lid region functioning is the bacterial mode retained by the *Plasmodium* species.

Although lacking the intrachain ATP-PPase-GATase interaction, the archaeal MDA is similar to the bacterial one (**Figure.3a**). The predicted archaeal GMPS complex with the GATase bound to the ATP-PPase subunit is similar to the unrotated *Plasmodium* GMPS^27^. Since archaeal GMPS FunFam have hydrophobic and aromatic residues conserved on SDPs 222, 370 and 415 similar to the *Plasmodium* GMPS FunFam (**Table.3**), the bound archaeal GATase subunit may undergo further rotation and activation like the*Plasmodium*GMPS^27^. The key difference between archaeal GMPS ATP-PPase domain and the other GMPS is the SDP residue R/W373. All the rest of the GMPS FunFams conserve non-aromatic hydrophobic residues at this position (**Table.3**). The function of this archaeal SDP373 residue has not been investigated experimentally. Since only archaeal GMPS performs transient GATase subunit binding to the ATP-PPase subunit, further investigation of this residue may clarify the mechanism of the archaeal GMPS complex formation.

Eukaryotic GMPSs have a unique MDA compared with prokaryotic GMPSs. Their homodimer has a two-layer pyramid shape, with the two ATP-PPase and four dimerization domains on the bottom and the two GATase domains on the top (**Figure.3b, c**). Thus, the ATP-PPase domains are interacting with a dimerization domain of the other protomer in contrast to the prokaryotic GMPSs which interact with the GATase domains. The Lid region is therefore not able to interact with the GATase domain (**Figure.3b, c**). Activity assays together with centrifugation studies have revealed that eukaryotic GMPSs function as monomers^54,55^, which is also in contrast to the prokaryotic homodimers^6,7,47^. It has been hypothesized that eukaryotic GMPS transiently dimerize for substrate XMP binding, and subsequently dissociate to catalyse XMP-adenylation and GMP-synthesis^28^. The additional eukaryotic dimerization domain may be stabilising the GMPS monomer. In addition, it is possible that XMP-adenylation and the Lid region movements regulate the dimer dissociation and GATase domain relocation. This hypothesis is supported by eukaryotic GATase domain activity assays. The eukaryotic GATase domains were found to be activated only when all ligands for XMP adenylation (XMP, ATP, Mg^2+^) are bound to the ATP-PPase catalytic site^54^.

We examined whether the eukaryotic FunFam specific SDP residues could promote dimer dissociation by interacting with the GATase domain and contributing to GATase domain relocation. There are 4 SDP residues, SDP394-396 and 398, which sequentially follow the catalytic Lid motif (388-393), and for which the eukaryotic FunFam has a unique conservation pattern (**Table.3**). Therefore, these SDP residues may be involved, but there is currently no experimental information available on the dynamics and mechanisms of this process. The eukaryotic SDPs 222, 370, 394-396, and 415 have charged residues conserved only in the eukaryotic GMPS whilst the prokaryotic homologs possess hydrophobic residues. Further investigation of whether these charged residues interact with the GATase domain SDPs may provide new insights into the mechanisms by which changes in the eukaryotic oligomers occur.

### The Lid region potentially becomes structured to form an ammonia tunnel for all GMPS

The exact mechanism by which the GMPS transport the NH_3_ produced by the GATase domain to the ATP-PPase domain catalytic site is still uncertain. However, it has been proposed by various studies that the intrinsically disordered Lid region located on the interface between these two catalytic domains can facilitate the formation of an NH3-tunnel^7,16,46,48^. Since NH_3_ transportation is a common function of all GMPS, fully conserved residues within the Lid region may be not only critical for catalysing XMP adenylation^7,48^, but also binding or interacting with the NH_3_ molecule to transport it to the GMPS active site. Interestingly, the GMPS specific Lid motif IK(T/S)HHN contains two His residues and the conservation of such His pairs lining the wall of an NH_3_-tunnel is a known feature of the ammonia-transporting Amt/MEP protein family^56,57^. Hence, future experimental studies should further probe the role of the GMPS Lid motif to better understand the GMPS ammonia transportation mechanism.

### Characterising a GMPS Bacterial FunFam specific structural motif which distinguishes between eukaryotic and bacterial GMPS

Our analyses above revealed differential conservation of residues in the GMPS Lid region. SDPs 370, 414 and 415 were defined as structural motif residues. The residue conservation pattern is typically YLF and RFH in bacterial and eukaryotic GMPS FunFams respectively. In addition, a PBL motif residue D248^2^ conserved by all GMPS is also considered part of the structural functional motif. The specificity of the D248, Y370, L414 and F415 structural arrangement (DYLF motif) in bacterial GMPS SDPs was determined by annotating their occurrences in a large set of 3,637 high quality AF2 predicted HUP domains using the Amino acid pattern Search for Substructures and Motifs (ASSAM) webserver^58^. ASSAM matched 54/56 bacterial and *Plasmodium* FunFam members to the DYLF motif (**Figure.4**) with no eukaryotic matches. The two unmatched bacterial GMPS have a Tyr instead of a Phe at SDP415. Although we predict this as a conserved SDP site, these residues have similar physicochemical properties, and the mutation has been tolerated.

**Figure.4.**
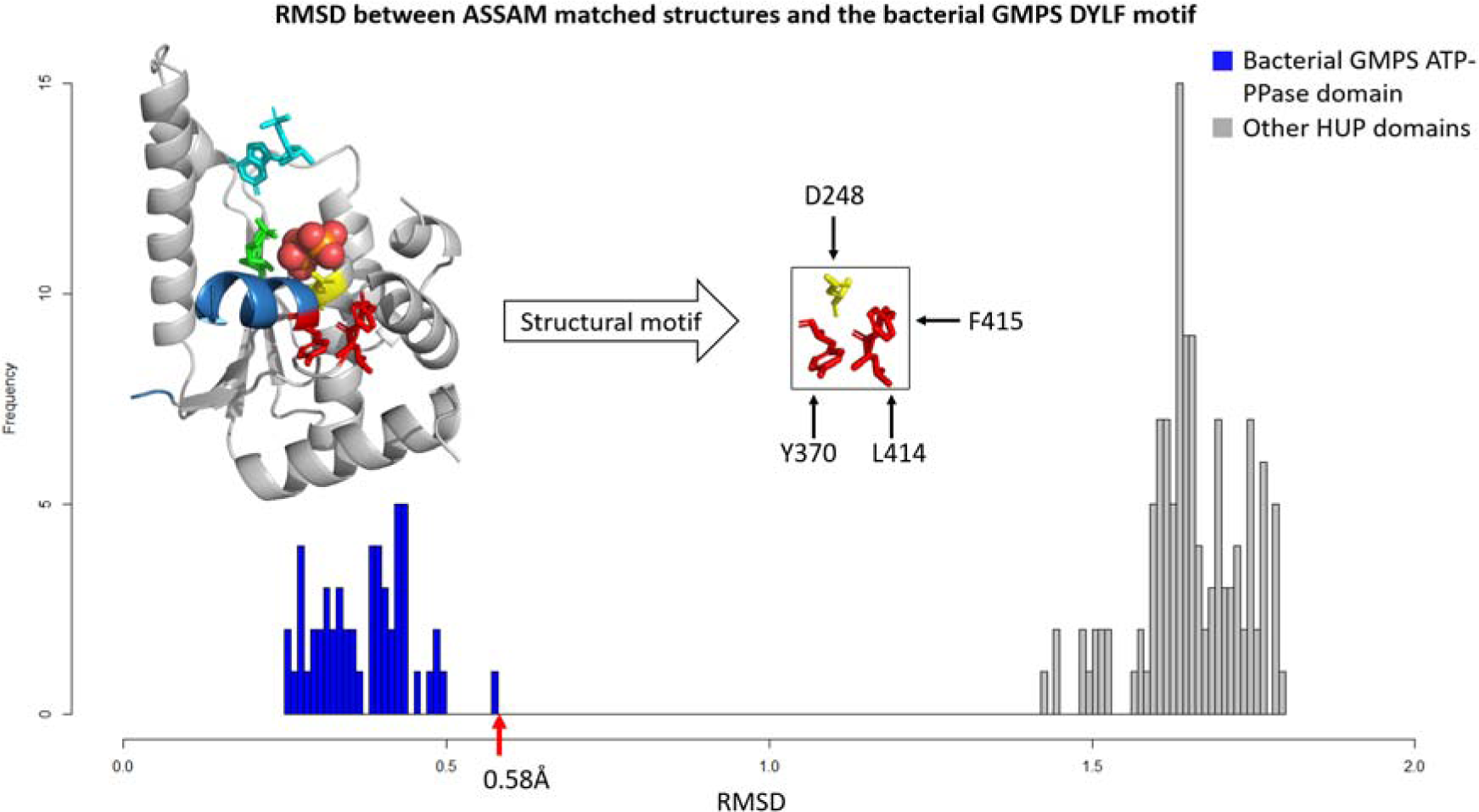
Matches obtained by searching the bacterial GMPS-specific structural motif against the HUP dataset of structures using ASSAM. PDB: 1gpm. Colour scheme: Blue-Lid region; Yellow-Phosphate Binding Loop motif; Red-GMPS SDP residues; Cyan-XMP; Green-AMP; Red and orange sphere-PPi.

A subsequent ASSAM search using the putative bacterial specific Gln-NADS motif against the whole PDB and AF2-predicted human structures did not retrieve any significant matches in human proteins. Furthermore, the CavityPlus Webserver^59,60^ analysis of the bacterial FunFam representative Escherichia coli GMPS (PDB: 1gpm) predicted a highly druggable cavity on the ATP-PPase domain (**Supplementary. Figure2**) containing the DYLF structural motif suggesting that this motif identified by our computational analysis might facilitate anti-bacterial drug design making this domain a good drug target.

## Analysis of NADS FunFams and their function determining residues and structural regions

### Zebra3D identified a potential allosteric site specific to eukaryotic NADS

Unlike the GMPS, the substrate-binding loop of NADS exhibits a fully conserved structure (**Figure.2a**). However, Zebra3D analysis of the NADS ATP-PPase domains revealed a FunFam-specific SDR located away from the substrate-binding pocket (**Figure.5**). This region extends from the C-terminal of the α1-helix to the N-terminal of the β2-strand of the HUP common core (**Figure.5a**). Zebra3D subclassification of NADS based on this SDR resulted in a good match between the six Zebra3D subgroups and the six NADS FunFams (**Table.4**). However, there is an exception with the two bacterial NH_3_-dependent NADS FunFams (NH_3_-NADS). Their SDR simply comprises of the loop linking the α1 helix-β2 strand and therefore Zebra3D was unable to distinguish bacterial NH_3_-NADS structures as they are highly similar in this SDR resulting in one merged subgroup (**Table.4**).

**Figure.5.**
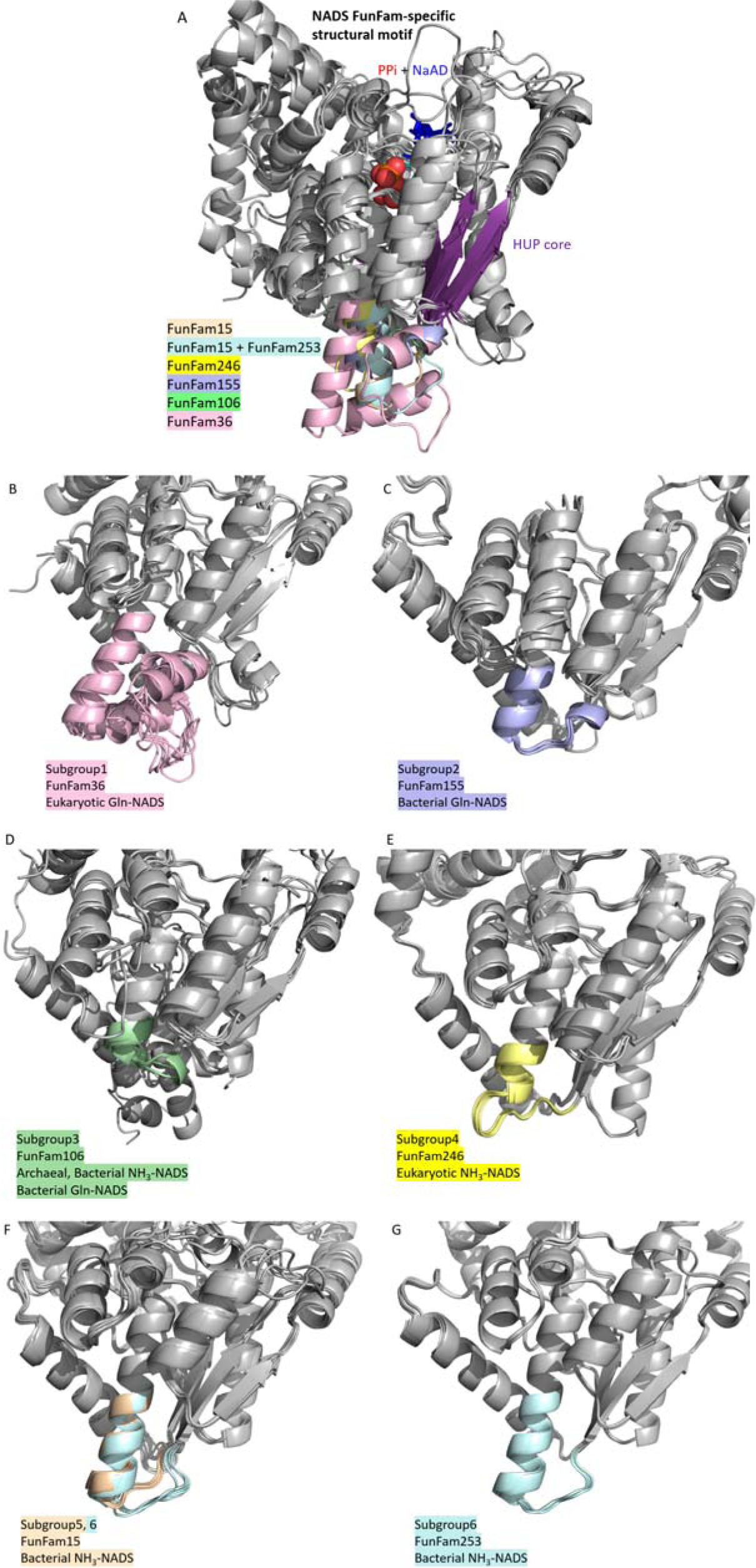
Zebra3D identified NADS FunFam specific SDR. (a) Superposition of the six representative structures from each NADS FunFam, SDR coloured according to Zebra3D sub-classification. (b-g) Superposition of NADS ATP-PPase domains from each FunFams.

**Table.4.**
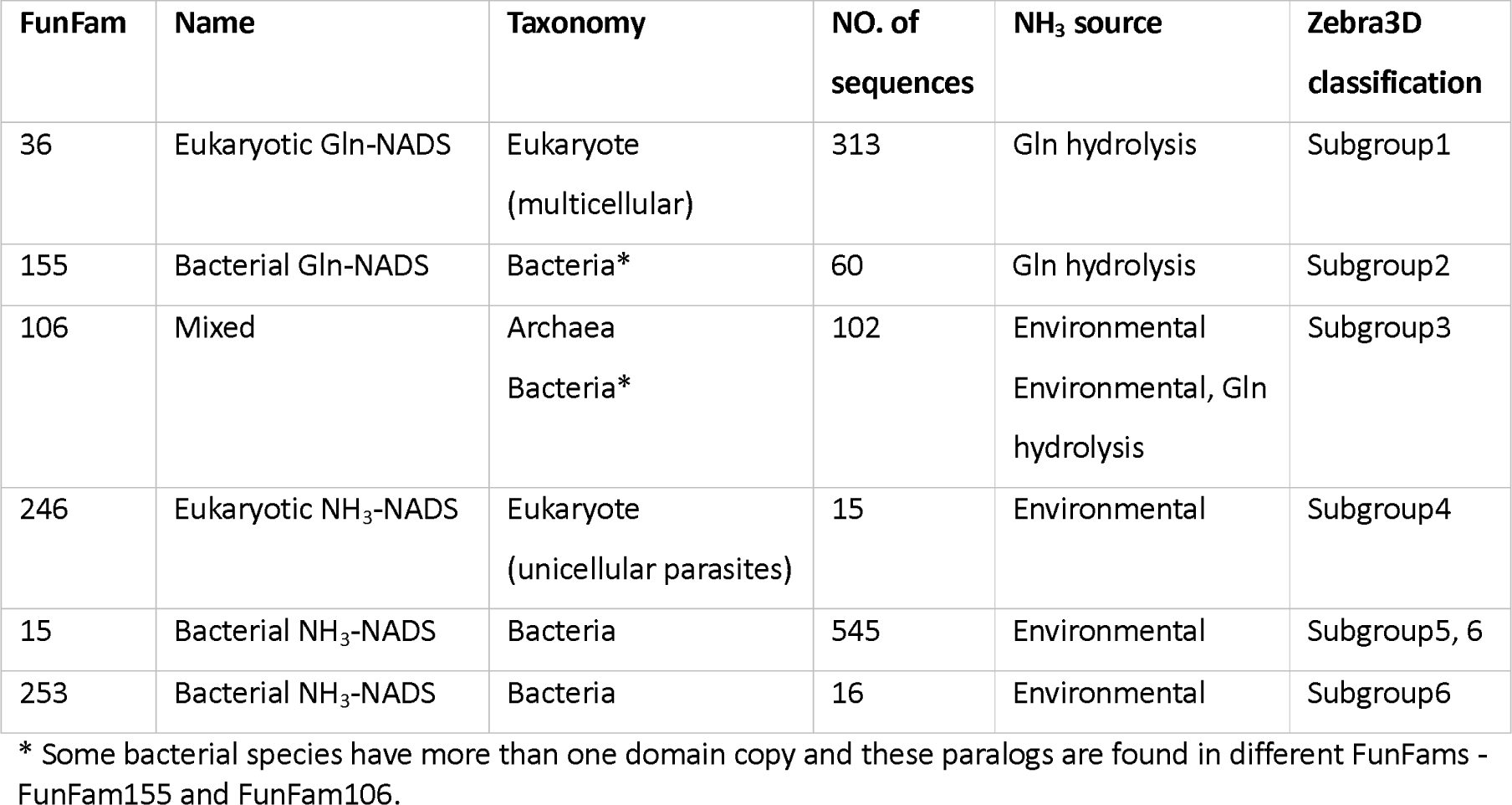
Zebra3D subclassification of NADS is highly comparable to FunFams subclassification.

| **Fun Fam** | **Name** | **Taxonomy** | **NO. of sequences** | **NH<sub>3</sub> source** | **Zebra3D classification** |
| --- | --- | --- | --- | --- | --- |
| 36 | Eukaryotic Gln-NADS | Eukaryote<br>(multicellular) | 313 | Gln hydrolysis | Subgroup1 |
| 155 | Bacterial Gln-NADS | Bacteria* | 60 | Gln hydrolysis | Subgroup2 |
| 106 | Mixed | Archaea<br>Bacteria* | 102 | Environmental<br>Environmental, Gln<br>hydrolysis | Subgroup3 |
| 246 | Eukaryotic NH<sub>3</sub>-NADS | Eukaryote<br>(unicellular parasites) | 15 | Environmental | Subgroup4 |
| 15 | Bacterial NH<sub>3</sub>-NADS | Bacteria | 545 | Environmental | Subgroup5, 6 |
| 253 | Bacterial NH<sub>3</sub>-NADS | Bacteria | 16 | Environmental | Subgroup6 |

\* Some bacterial species have more than one domain copy and these paralogs are found in different FunFams - FunFam155 and FunFam106.

The eukaryotic Gln-NADS FunFam specific SDR structure contains an additional short alpha-helical segments elements (**Figure.5a**). The role of these structural embellishments in the eukaryotic Gln-NADS α1-β2 loop has not been reported previously. The Protein Allosteric and Regulatory Sites (PARS) webserver^61^ predicted a potential allosteric site in eukaryotic Gln-NADS in which this SDR is involved (**Figure.6**). The presence of an allosteric site in eukaryotic Gln-NADS supports its distinct mechanism of regulation compared to bacterial Gln-NADS (details discussed below)^31,35^.

**Figure.6.**
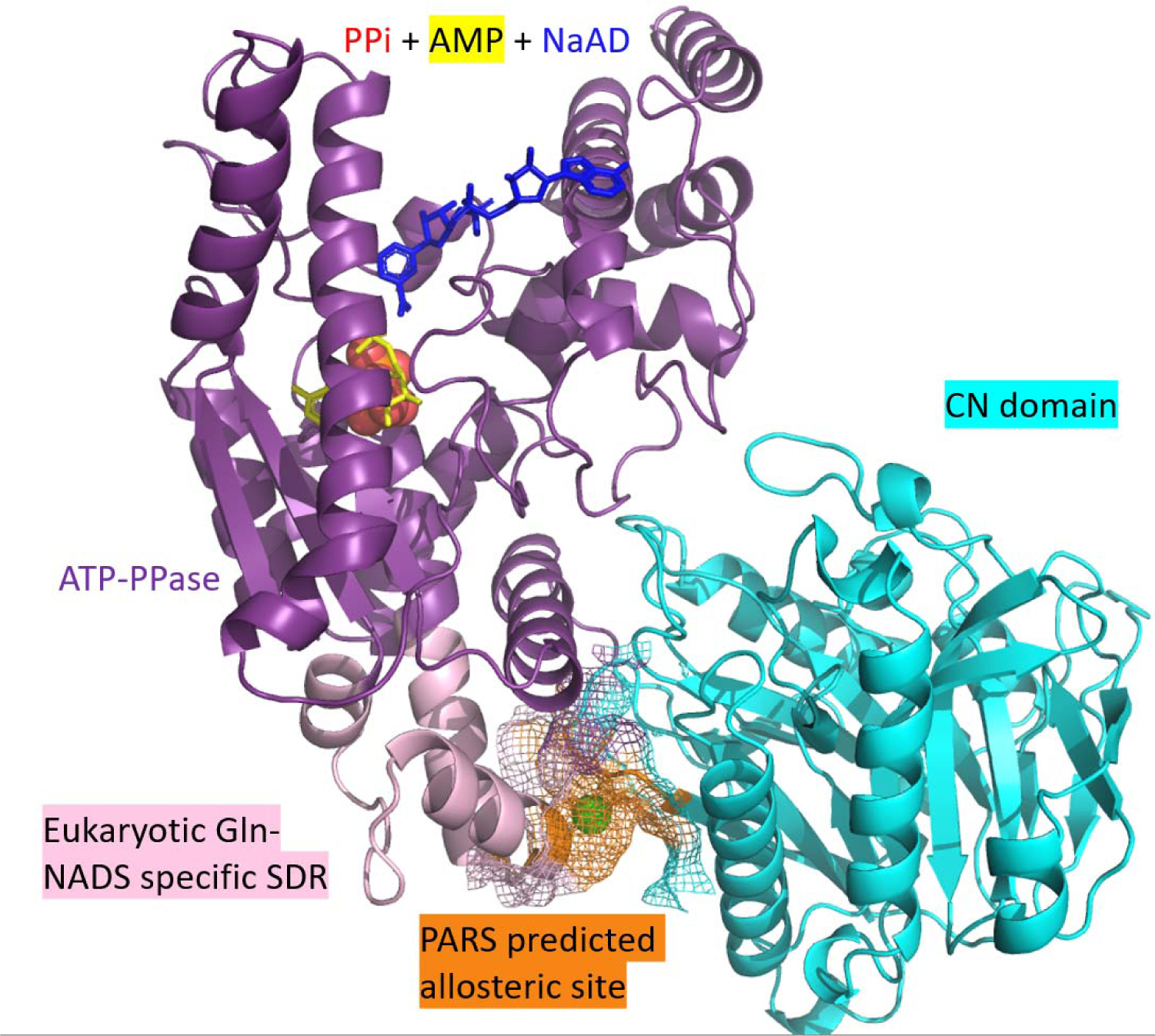
PARS predicted eukaryotic Gln-NADS specific allosteric site. PDB: 6ofb.

### Most of the NADS FunFam-specific SDPs are interacting with an ammonia tunnel

GroupSim analysis identified 10 Specificity Determining Positions (SDPs) with distinct residue-conservation patterns between the 5 NADS FunFams (**Table.4**). These positions are numbered according to the human Gln-NADS (PDB: 6ofb) (**Figure.7a, b; Table.5**). The locations of these SDPs can be assigned to three categories: (1) Part of or interacting with an ammonia tunnel (NH_3_-tunnel); (2) Interacting with an NADS characteristic functional loop known as the P2 loop^13,15^; (3) Adjacent to (∼5Å distance) to the NADS FunFam specific SDR.

**Figure.7.**
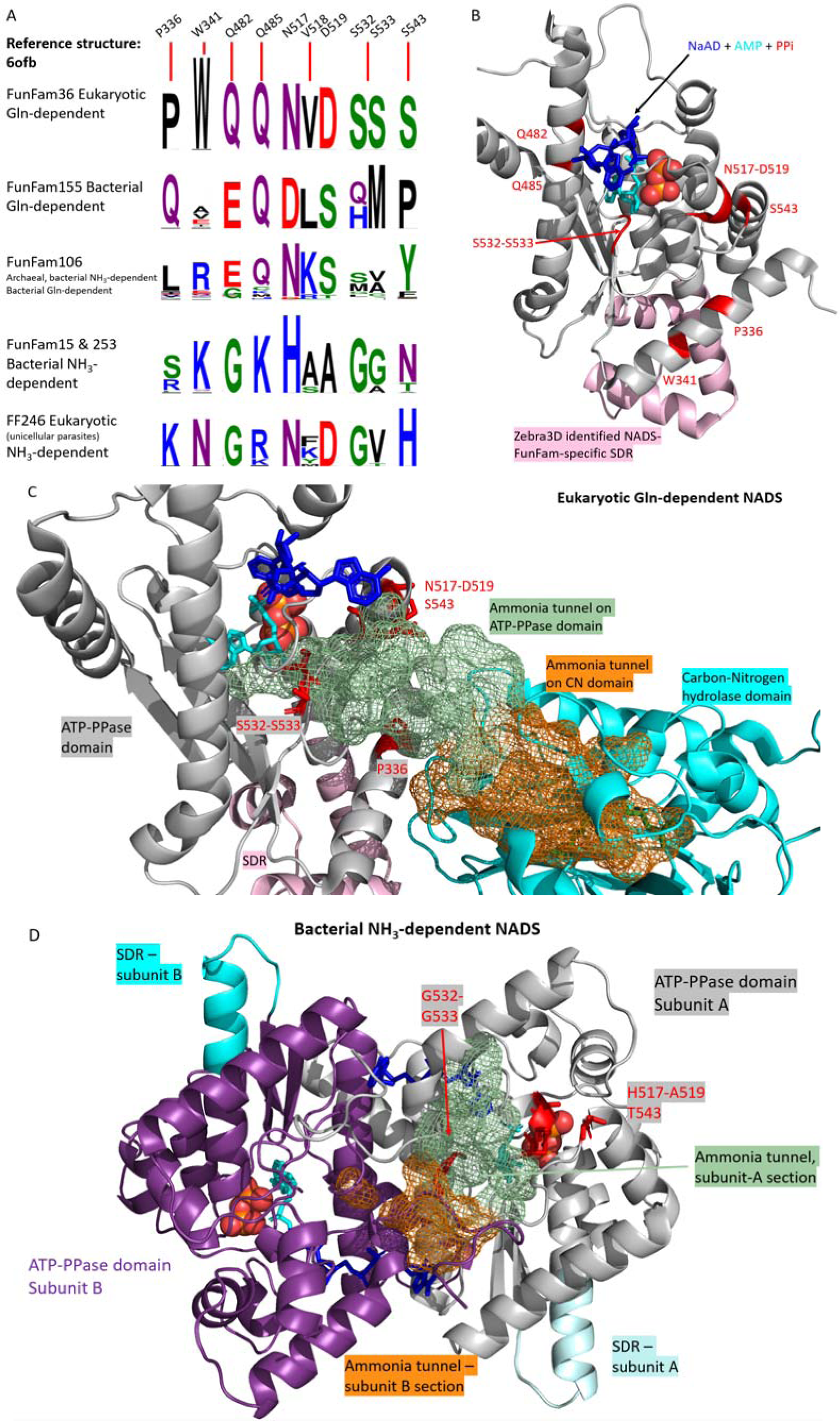
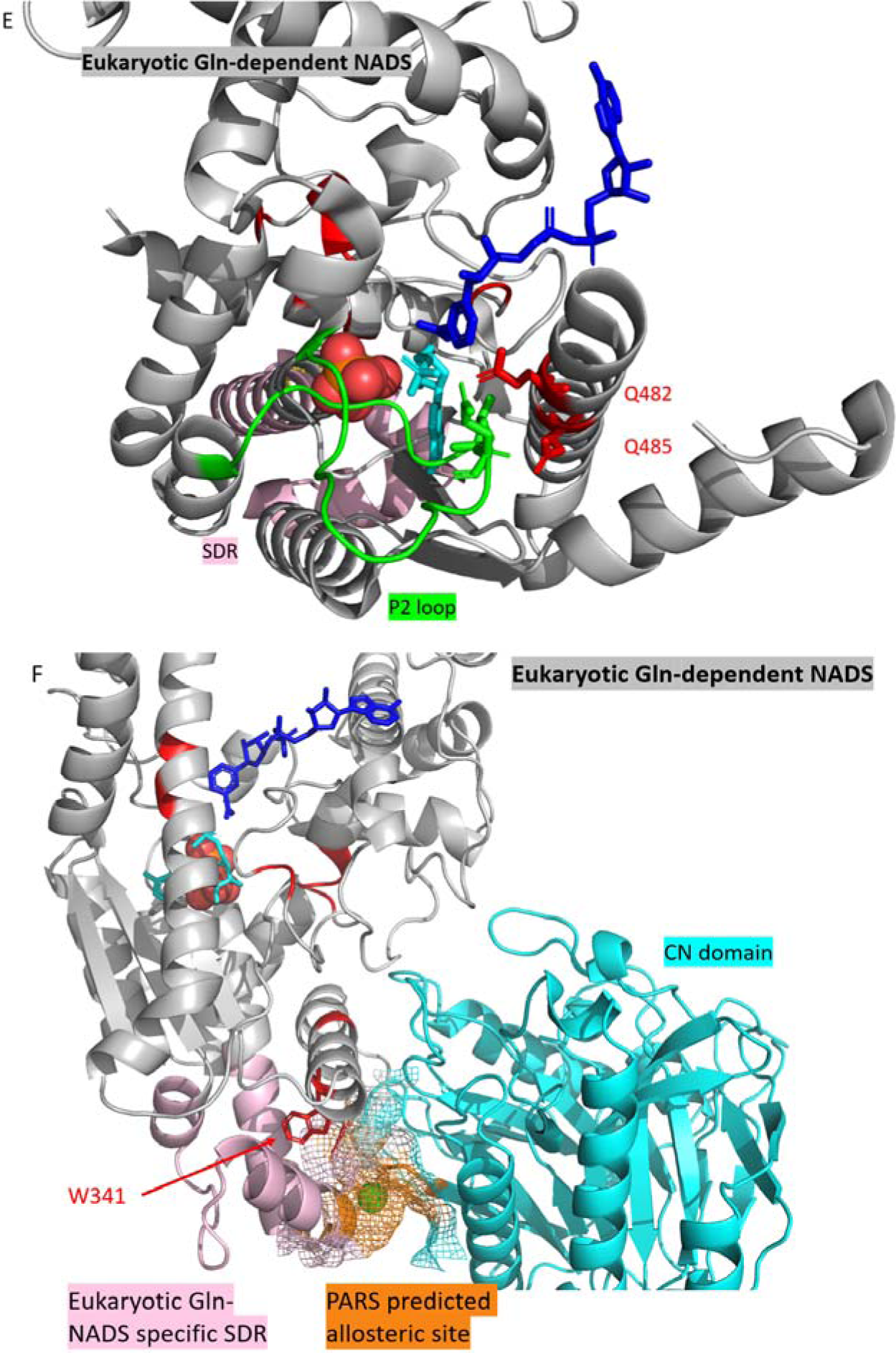
SDR and SDPs of NADS are found near to known functional sites. (a) GroupSim identified SDPs and their conservation patterns of each FunFam, figure generated by WebLogo3^51^. (b) SDR and SDPs mapped to human NADS (PDB: 6ofb). (c) NH_3_-tunnel (mesh representation) of human NADS with nearby SDR and SDP labelled. The tunnel residues are known from the literature^31^. (d) NH_4_^+^-tunnel (mesh representation) of bacterial NH_3_-NADS (PDB: 6kv3) The tunnel residues are known from the literature^36^. (e) P2 loop interacting with SDP residues 482 and 485. (f) SDP341 is adjacent to the FunFam specific SDR. Colour scheme for ligands: blue-NaAD; Cyan-AMP; Red-PPi.

**Table.5.**
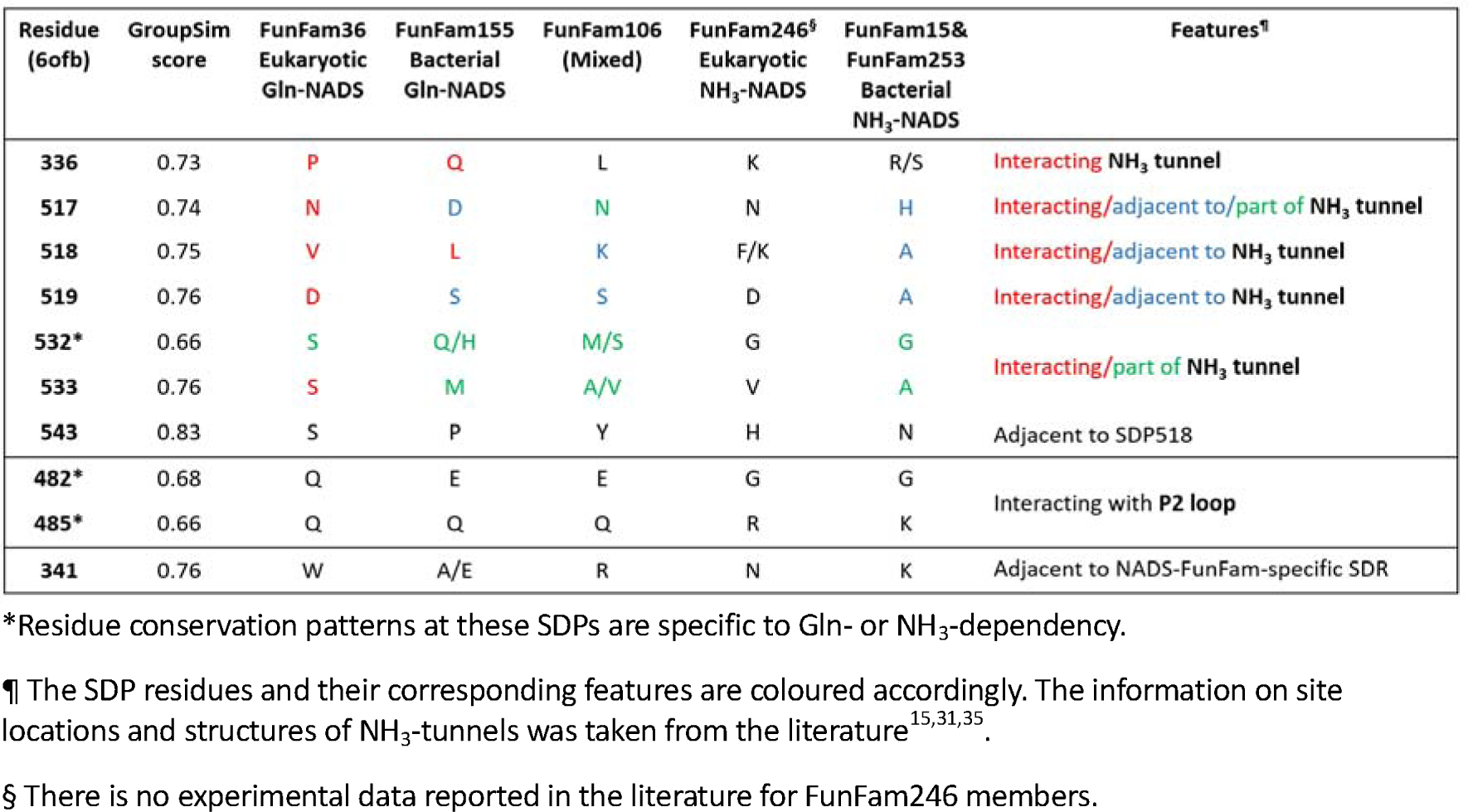
GroupSim identified Specificity Determining Positions across the four GMPS FunFams.

There are 7/10 SDPs associated with an NH_3_-tunnel which delivers NH_3_ to the active site of the NADS (**Table.5**). As with the co-operation between the GMPS ATP-PPase and GATase domains, Gln-NADS also recruit a glutamine-hydrolysing domain known as the CN domain. NH_3_ is produced by the CN domain hydrolysing Glutamine^15^. Existence of an NH_3_-transporting tunnel linking the CN and ATP-PPase domain catalytic sites located on the interface of these two domains has been predicted by the CAVER program in earlier studies of this subfamily^15,31,35,62^ (**Figure.7c, d**). The NH_3_-NADS are single domain enzymes with no CN domain, thus they have no interdomain tunnels. However, they have been proposed to transport ammonia/ammonium ion (NH_4_^+^) obtained from the environment from the binding site to the catalytic site, also *via* a tunnel^36^. Based on the molecular dynamics analysis by Sultana and Srivastava, this tunnel was deduced to be on the interface between two protomers of an NH_3_-NADS homodimer.

The remaining 2/10 SDPs were found on the other side of the NADS ATP-PPase domain catalytic pocket. Residues in these positions interact with the P2 loop (**Figure.6e**). The function of this loop is possibly associated with the NH_3_-tunnel in bacterial Gln-NADS^15^ (details in the following section). These SDPs have not been detected by previous studies.

The remaining SDP is about 5Å from the NADS SDR which has been predicted to be part of an allosteric site of the eukaryotic Gln-NADS FunFam (**Figure.6f**).

### Different Multidomain Architectures of the Gln-NADS FunFams explain differences in the SDP residues, the potential allosteric site and the associated functional mechanisms

All three Gln-NADS containing FunFams have varied MDAs. The Mixed FunFam bacterial Gln-NADS **(Table.4**) form homodimers with inter-subunit NH_3_-tunnels. Their NH_3_-tunnels link the catalytic site of the ATP-PPase domain from one monomer to the catalytic site of the CN domain of the other monomer of the dimer^35^ (**Figure.8a**). Bacterial FunFam Gln-NADS function as homo-octamers instead, and this octamer is a quadraplex of homodimers whose orientations are like the Mixed FunFam members^15,35^ (**Figure.8b**). Thus, the Bacterial FunFam members also possess inter-subunit NH_3_-tunnels but adopt a different oligomeric state. This resemblance suggests that Gln-NADS evolved from a deep-branching bacterial clade as a homodimer, to the more recent bacterial Gln-NADS that function as homo-octamers^35^. In contrast, the Eukaryotic FunFam Gln-NADS function as homo-octamers but form intra-subunit NH_3_-tunnels, which probably evolved later than the bacterial tunnels^35^ (**Figure.8c, d**). The PARS predicted allosteric site of eukaryotic Gln-NADS is found on the interface between the ATP-PPase and CN domains of the same subunit as well as the NH3-tunnel (**Figure.8e**).

**Figure.8.**
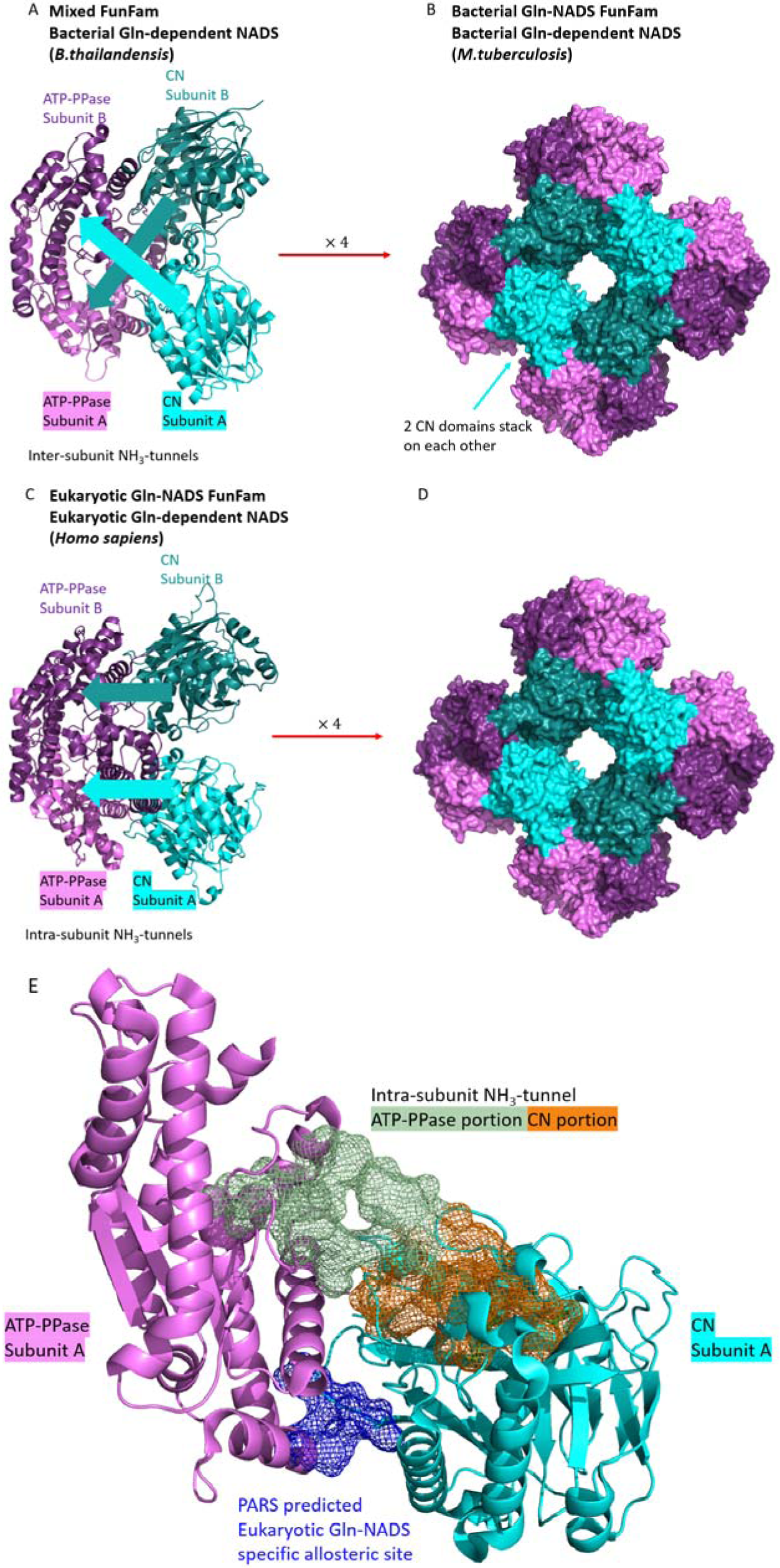
MDA variation across Gln-NADS FunFams. (a, b) Bacterial Gln-NADS from FunFam106 and FunFam155 respectively (PDB: 4f4h, 3syt). (c, d) Eukaryotic Gln-NADS from FunFam36 (PDB: 6ofb). (e) CAVER predicted eukaryotic Gln-NADS NH_3_-tunnel^31,62^ and PARS predicted allosteric sites^61^ mapped onto a eukaryotic Gln-NADS subunit.

These varied MDAs, explain differences in the seven residue NH_3_-tunnel associated SDPs of the three Gln-NADS FunFams (**Table.5**), although all the NH_3_-tunnels can regulate NH_3_ transportation by their relaxation and restriction^15,31,35^. Typically, there are multiple constriction sites across the NH_3_-tunnel where the radius of the tunnel is significantly narrower than the remaining parts. Relaxation of these sites is necessary for the NH_3_ to pass through^15^. All Gln-NADS have been observed to have a constriction site near the ATP-PPase catalytic site, acting as a gate^15,31,35^. The seven SDPs encircle this constriction site (**Figure.7c**). These SDP residues had not been detected by previous studies and we suggest that they may have a role in controlling the relaxation of this constriction site by interacting with the tunnel-wall residues.

The inter-subunit nature of ammonia transfer in Bacterial FunFam Gln-NADS probably resulted in its unique regulatory mechanism which relies on the P2 loop movement^15,31^. The P2 loop is an intrinsically disordered loop and conformational change in this loop on M. *tuberculosis* NADS (MtNADS) has been observed to activate the Gln-hydrolysis of a neighbouring CN domain in a different subunit^15^. Since Bacterial Gln-NADS FunFam members form inter-subunit NH3-tunnels, two of the eight protomers are involved in this tunnel formation, and the P2 loops on their ATP-PPase domains regulate the third and fourth CN domains instead of their own^15^. Therefore, the Mixed-FunFam members which form homodimers probably do not perform such regulation, and the P2 loop of the eukaryotic Gln-NADS was found to have no function in regulating CN domain activity^31^.

Instead of P2 loop regulation, the eukaryotic Gln-NADS probably evolved to a unique regulatory mechanism relying on the allosteric site. This is supported by experimental assays of activity of the MtNADS^15^ and human Gln-NADS^31^ showing that bacterial Gln-NADS strongly prefers Gln over environmental NH_4_^+^ as its ammonia source^15,63^, while human Gln-NADS has no preference^31^. However, if the eukaryotic Gln-NADS really has no preference between these ammonia sources, the ATP-PPase and CN domains could simultaneously bind NH_4_^+^ and Gln to their catalytic sites. This would be a waste of Gln when the CN domain generates an NH_3_ but the ATP-PPase domain is using an NH_4_^+^ for the second step of NADS production. If the NH_3_ produced by the CN domain is not consumed by ATP-PPase domain, it would be released to the cell environment^35^. This is contradictory to the concept of ATP-PPase-CN domain synergism. Therefore, it is possible that the eukaryotic Gln-NADS regulate their affinity towards Gln via their unique allosteric site (**Figure.8e**).

### Structural motif consisting of SDPs adjacent to the Gln-NADS ammonia tunnel can be used to distinguish human and pathogenic bacterial NADS

Human and several pathogenic bacteria including *M. tuberculosis* express Gln - NADS^15,31^. Hence, we focused on identifying structural motifs that can distinguish between the eukaryotic and bacterial Gln-NADS FunFams. The structural motif was defined as SDP residues 336, 517-519 and 543, which are in proximity to each other and to the NH_3_-tunnel (**Figure.7c, Supplementary. Figure3**). The residue conservation patterns are QDLSP and PNVDS for bacterial and eukaryotic Gln-NADS respectively.

For the bacterial QDLSP structural motif, ASSAM searches matched all five AF2 Bacterial FunFam Gln-NADS ATP-PPase domains (**Figure.9**), with no other HUPs matched significantly. Similarly, searching the eukaryotic PNVDS motif resulted in matching only eukaryotic Gln-NADS (**Supplementary. Figure4**). ASSAM queries using the bacterial QDLSP motif against the whole PDB and AF2-predicted human proteins did not return any significant matches to human proteins^37^. CavityPlus^59,60^ predicted a highly druggable pocket on the MtNADS (PDB: 3syt) where the QDLSP motif is buried deep inside (**Supplementary. Figure5**). Therefore, this structural motif could be further investigated for anti-bacterial drug development.

**Figure.9.**
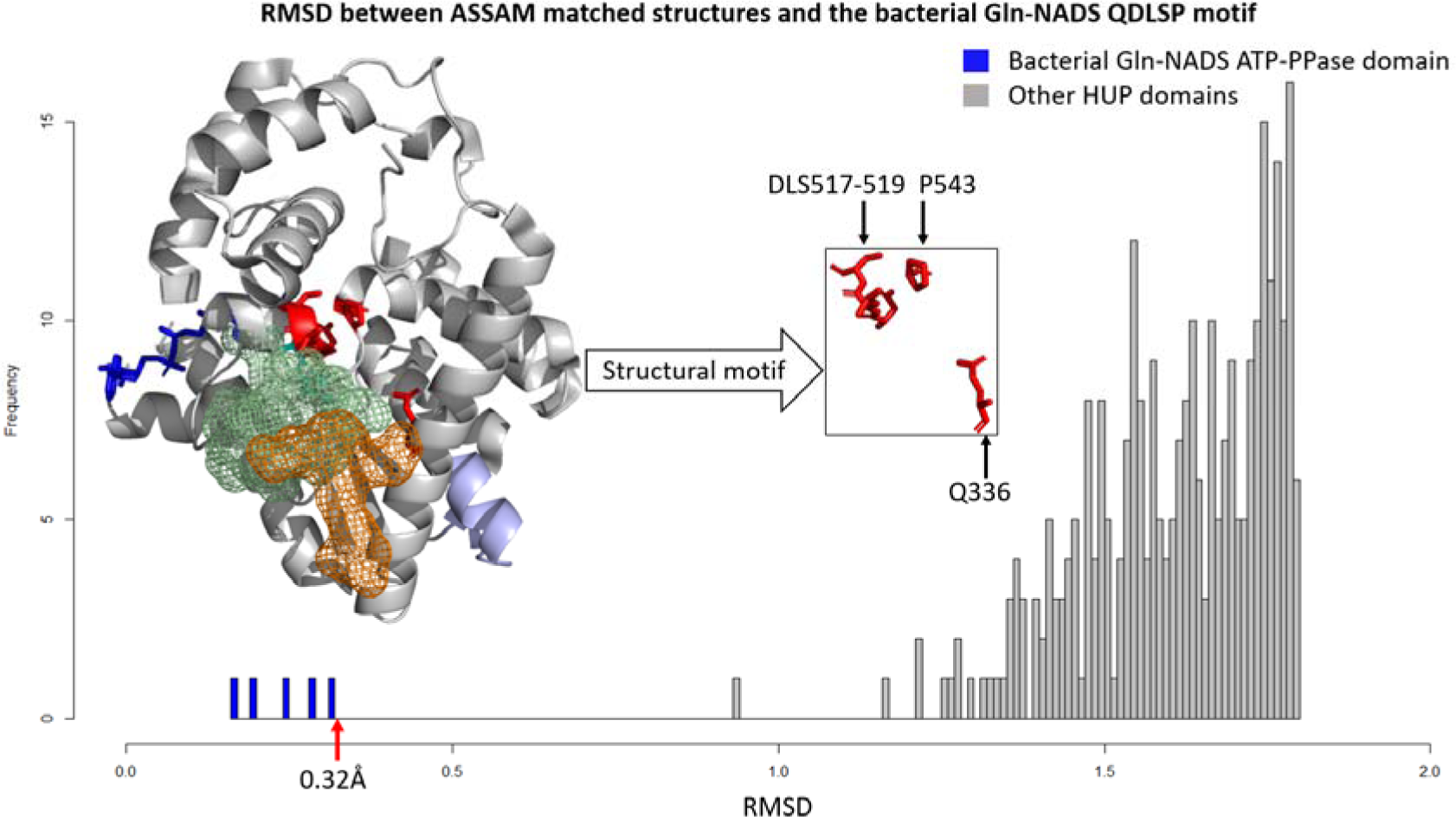
Results from searching the bacterial Gln-NADS specific structural motif against a dataset of HUP structures using ASSAM. PDB: 3syt. Colour scheme: Light purple-NADS FunFam-specific SDR; Blue-NaAD; Red-SDP residues; Light green and orange mesh-ATP-PPase- and CN-domain portion of NH_3_-tunnel^15^.

### Convergent evolution of GMPS and NADS Gln-dependency

Structural and sequence comparisons of GMPS and NADS suggest convergent evolution of their Gln-dependency. While their ATP-PPase domains both belong to the HUP superfamily, their Gln-hydrolysing domains belong to two separate superfamilies in the CATH database, the GATase (CATH: 3.40.50.880) and the CN (CATH: 3.60.110.10). Superposition of a GMPS GATase domain (PDB:1gpm) and a NADS CN domain (PDB: 4f4h) revealed low structural similarity (TM-score = 0.39) that suggests the lack of any evolutionary relationship^64^. Therefore, their similar dependency on Gln-hydrolysis as an NH_3_ source is probably the result of convergent evolution. A phylogenetic tree of ATP-PPase domains was constructed to obtain more details of this evolutionary history.

The phylogenetic tree was inferred (**Figure.10**) by aligning ATP-PPase sequences from 36 FunFams (**Supplementary. Table1**), including the four GMPS and six NADS. There are two interesting features of this tree. Firstly, the GMPS ATP-PPase domains form the deepest branch of ATP-PPase, with the NADS more recently emerged. Secondly, although the NADS ATP-PPase domains form a separate clade on this phylogenetic tree, there are clearly two distinct subclades corresponding to NH_3_- and Gln-NADS. As some of the NH_3_-NADS branches are significantly shorter than for Gln-NADS, this tree suggests that NH_3_-NADS emerged earlier than Gln-NADS, in agreement with previous genetic studies of NADS^65^.

**Figure.10.**
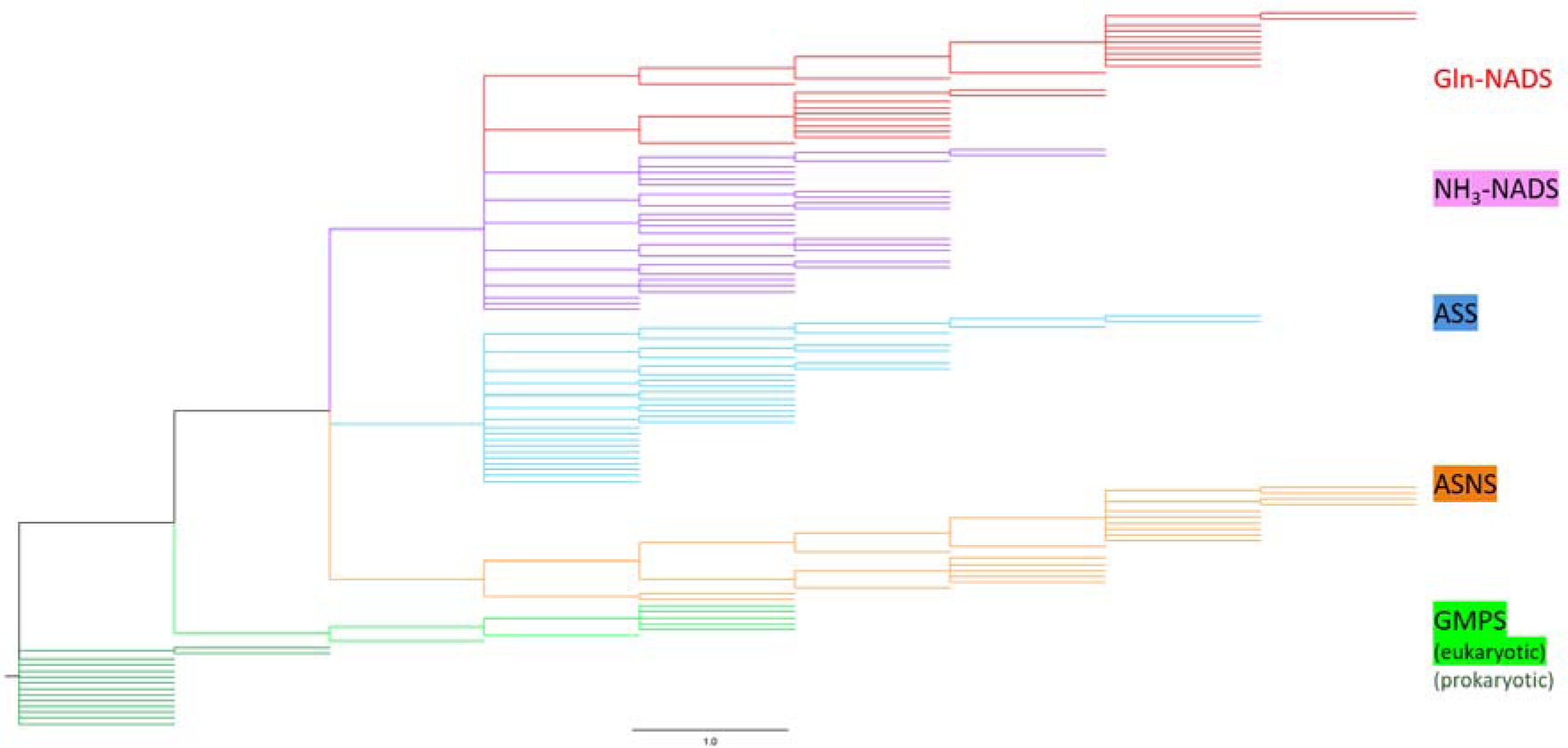
Phylogenetic tree of ATP-PPase domains across 36 FunFams and four substrate-specificity groups. Colours are indicated by branch labels. Eukaryotic GMPS refers to FunFam44 members only; Prokaryotic GMPS includes bacterial, archaeal and Plasmodium GMPS.

Based on the tree, we hypothesize that the emergence of CN domains occurred later than the GATase domain. As GMP and NAD are both life-essential molecules^29,30^, the ATP-PPase common ancestor had probably diverged to GMP- and NAD-synthesizing domains before the Last Universal Common Ancestor (LUCA) split into different life forms^1^. GATase might be another pre-LUCA enzyme which evolved to cooperate with GMPS ATP-PPase domains. In contrast, presumably no CN domain emerged early to participate in NAD synthase.

Our study was focused on the differences between GMPS and NADS ATP-PPase domain structures and sequences. Further computational analyses of GATase and CN superfamilies could provide insights into the evolution of the GMPS and NADS Glutamine-dependency. Our protocol could also be applied to other ATP-PPase domains from Asparagine Synthase and Argininosuccinate Synthase to extend our understanding of ATP-PPase domain evolutionary history.

## Conclusions

ATP-Pyrophosphatases from the large and diverse HUP domain superfamily exhibit considerable plasticity in substrate binding and have evolved different multidomain architectures. ATPases having different substrate-specificity groups cooperate with different Glutamine-hydrolysing domains to obtain their nitrogen-containing substrates. Within a substrate-specificity group, different domains of life have their specific mechanisms of domain synergism between ATP-PPase and Gln-hydrolysing domains. Our computational analysis of two substrate-specificity groups, the GMPS and NADS ATP-PPase domains, compared their structures and sequences and comprehensively characterized the specific features distinguishing them. The computational results not only aligned well with experimental results, but also discovered additional key residues and potential allosteric sites previously uncharacterised. In addition, structural motifs specific to different taxonomic groups were identified by our computational protocol. Understanding these differences could contribute to the development of anti-bacterial drugs targeting the bacterial life-essential enzymes GMPS and NADS.

## Acknowledgement

N.S acknowledges funding from the Biotechnology and Biological Sciences Research Council (grant code: BB/V014722/1). V.M acknowledges funding from Welcome Trust grant [221327/Z/20/Z]. M.F.R acknowledges funding from Ministry of Higher Education Malaysia Translational Research Grant Scheme (TRGS/1/2022/UKM/01/9/1).

## STAR★Methods

### Key Resource Table

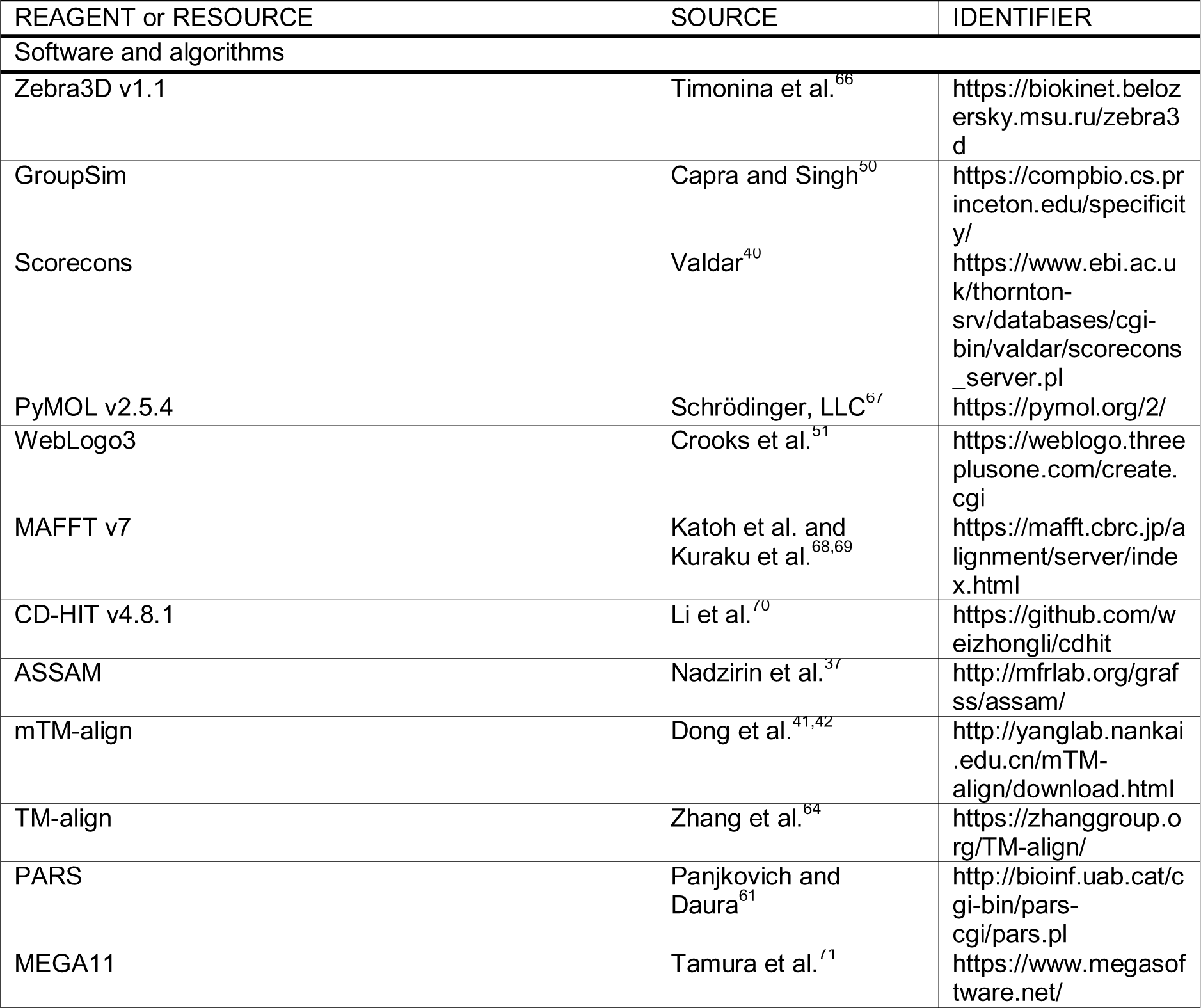
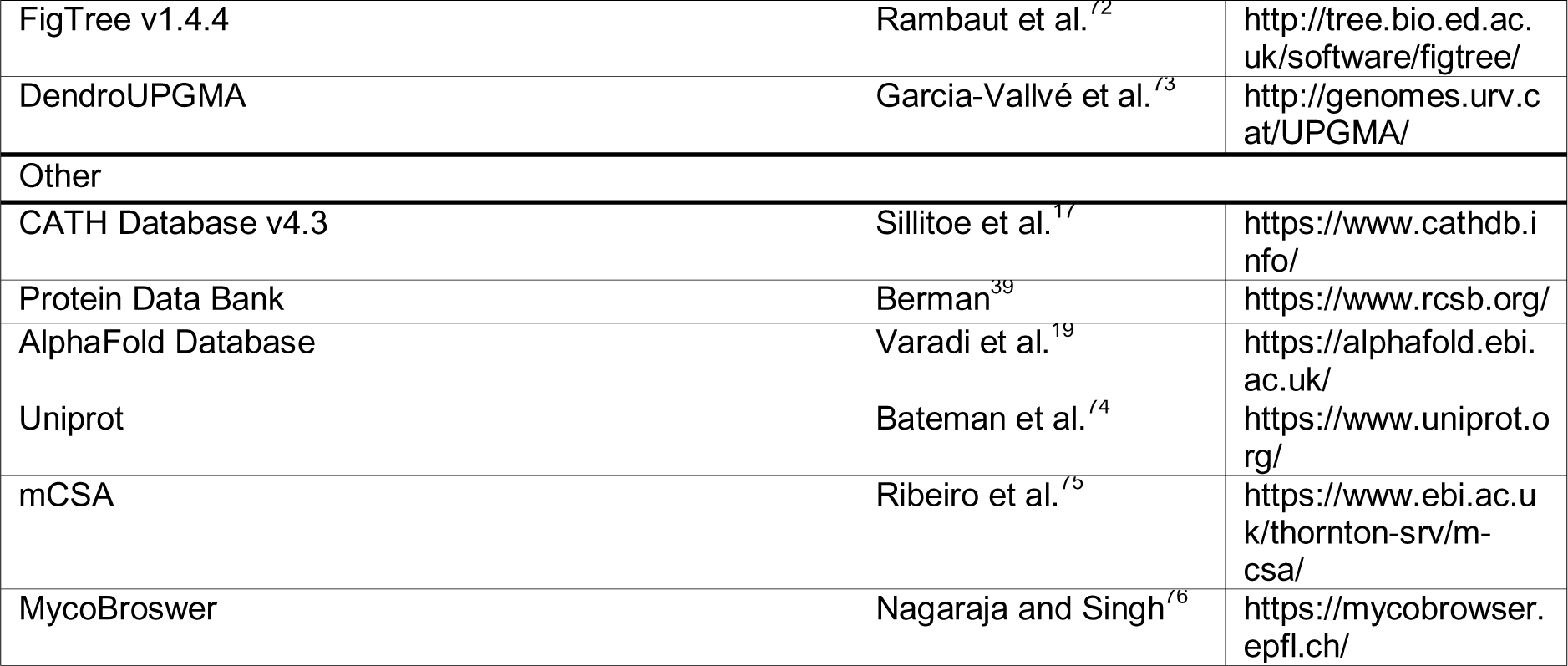

### Resource Availability

#### Lead contact

Further information and requests for resources should be directed to the lead contact, Christine Orengo.

#### Data and code availability

The dataset of AF2-predicted HUP domains with overall pLDDT > 90 are available at https://doi.org/10.5281/zenodo.8346482.

The analyses reported in this paper did not exploit novel algorithms and therefore we do not report any original code. All algorithms used have been cited.

## Method details

### CATH database and Functional Families

Our study focused on the HUP superfamily of CATH database Version 4.3 (CATH:3.40.50.620)^17^. There are 41,313 sequences belonging to HUP superfamily, where 1,685 of them have experimentally resolved structures deposited in PDB (Protein Data Bank)^39^.

Sequences within the superfamily are classified into functional subfamilies by first applying a hierarchical agglomerative clustering method (GeMMA)^22^ that progressively clusters sequences according to their similarity (as determined initially by pairwise sequence comparisons and subsequently by HMM-HMM comparisons between sub-groups). This is used to generate a tree of relationships which is then cut into functional groups using the FunFHMMer algorithm^77^. FunFHMMer segregates groups of functionally distinct relatives (i.e. FunFam) by identifying differentially conserved residues between them. These are Specificity Determining Positions (SDPs) specific to the individual FunFams and typically associated with functionally important residues e.g. catalytic, binding or interface residues^50^ (see also details in section **highly conserved residues by Scorecons** and **Identifying Specificity Determining Positions by GroupSim**). FunFams were annotated with Gene Ontology terms from UniProt-GOA automatically^23,38,77^.

### AlphaFold2 predictions of HUP superfamily members

AF2 models of HUP domains were extracted from the AlphaFold Database^18,19^. The raw protein models were chopped into domains via a Python pipeline based on pdb-selres module from pdb-tools^78^ specifically built for this purpose, and were assigned into the HUP CATH superfamily^20^.

A non-redundant set of HUP sequences was generated by CD-HIT algorithm^70^ clustering the sequences at 90% identity and then selecting a single representative from each cluster.

Multiple criteria were applied to the obtained HUP domain models to identify high-quality models. Firstly, for AF2 prediction quality measurement the predicted local-distance difference test (pLDDT) was used^18^. We averaged the pLDDT scores of all residues within a predicted HUP domain and removed the models with <70 average pLDDT^20^.

Secondly, models were removed if they contained a high proportion of residues not in a secondary structure^20^, as calculated by the Dictionary of Protein Secondary Structure (DSSP) algorithm^79^. Models with over 65% residues not forming secondary structures were excluded from the dataset^20^.

Thirdly the globularity of the predicted HUP domains was considered. Both the packing density and surface to volume ratio were calculated for each domain models. Packing density was calculated by number of residues in proximity (< 5Å distance) to a hydrophobic residue, and this number is averaged across all hydrophobic residues within a domain to represent its packing density^20^. This was done using the python package Bio.PDB^80^. The solvent excluded surface (SES) area and volume of each HUP model were obtained by PyMOL MSMS plugin^67^. The smaller the SES/volume ratio, the more globular a domain is^20^. The threshold of these two measurements were defined by applying these methods to 61,328 curated experimental CATH domain structures from all superfamilies. The density and SES/volume ratio of the top 95% domain of this curated dataset are 9.75 and 0.494 respectively. The AF2 HUP domain models with globularity below these thresholds were discarded^20^. In total 3,637 non-redundant high-quality AF2 predicted HUP domains were retained for this study.

### Data collection

HUP FunFams associated with ATP-PPase functions were identified using the CATH FunFHMMer webserver^21^ and verified manually. We followed the definition of ATP-PPase of the previous HUP superfamily studies^1,2,10,11^, identifying ATP-PPases by their Phosphate Binding Loop motif SGGxDS and their enzymatic functions. We started with 12 ATP-PPases domain structures identified by a previous HUP superfamily subclassification study available in PDB^10^.

The CATH FunFHMMer webserver was used to search the sequences of these 12 structures against the CATH database. Matched FunFams with multiple sequence alignments showing fully conserved SGGxDS motifs and an Enzyme Commission (EC) number referring to an ATP-PPase reaction were selected manually as ATP-PPase FunFams.

Furthermore, a threshold on the Diversity of Position Score (DOPS) above 70 was applied to filter out less informative FunFams. DOPS is a measurement of the diversity of the member sequences, provided by the Scorecons algorithm^40^. High DOPS of a FunFam suggests its members originate from various taxonomic groups instead of from a few closely related species. This means that highly conserved residues of high-DOPS FunFams are likely to be associated with functional site^21^. For FunFams with high DOPS but no experimental structure, AF2 predicted models were collected. Our analysis focused on 16 high-DOPS FunFams and 2 FunFams with low DOPS but have at least one experimental structure. These 18 out of 39 ATP-PPase FunFams consists of 85% of the 6,168 ATP-PPase sequences.

### Multiple Structural Alignment

Multiple structural alignments were constructed for non-redundant representatives of the ATP-PPases by the program mTM-align^41,42^. For FunFams with multiple experimental structures, models obtained from the same species were eliminated to reduce redundancy. For FunFams having none or less than three experimental structures but having a high DOPS, up to three AF2 predicted models were recruited to enrich the number of structures examined and to allow subsequent Zebra3D analysis.

In the outputs of mTM-align, the structures are superposed as a single PDB file, and the superposition-guided sequence alignment is available directly as a FASTA file. In addition, there is a pairwise similarity matrix of the input structures. To visualize the structural relationship between FunFams, we input the similarity matrix to the DendroUPGMA webserver^73^ to generate a dendrogram (http://genomes.urv.cat/UPGMA/).

### Local structural comparison by Zebra3D

The mTM-align-superposed domain structures were used as input of the Zebra3D program^66^, which identifies regions of the ATP-PPase domains which can be conserved in particular subsets of the input structures. These Specificity Determining Regions (SDR) are therefore used to identify subgroups which show conservation of 3D structure within a subgroup but vary between subgroups^66^.

The algorithm of Zebra3D works in three steps. Firstly, the structural-alignment guided sequence alignment of the ATP-PPase sequences is analysed to identify a common core region. Continuous positions on this alignment with high residue conservation are considered as the core region. The remaining more variable regions are automatically compared by superposing their C_α_ coordinates to detect if they are differentially conserved in some subsets. RMSD between each pair of input structures at the variable regions are calculated, resulting in a distance matrix. Finally, a machine-learning method, by default HDBSCAN^81^, is applied to cluster the input structures into subgroups according to the matrix^66^.

### Identification of allosteric sites by PARS

For some Zebra3D identified significant variable regions that are not part of previously known ATP-PPase functional sites, the possibility of them being an allosteric ligand-binding site was considered. The Protein Allosteric and Regulatory Site (PARS) webserver was applied to examine such possibilities^61^.

### Identifying highly conserved residues by Scorecons

Scorecons^40^ calculates the residue conservation score of each column of the input Multiple Sequence Alignments (MSA). For the ATP-PPases, default parameter settings of Scorecons were applied.

We manually collected the sequences of FunFams having GO and/or EC annotations for GMPS or NADS and then combined and re-aligned them using MAFFT^68^. Scorecons measured the level of conservation of this combined MSA and of individual FunFams. For the alignments with a DOPS greater than 70 we calculated the Scorecons value of the residues. The threshold of a high conservation score was defined as 0.7. The positions with highly conserved residues were proposed to be critical for the function indicated by their shared GO annotation.

### Identifying Specificity Determining Positions by GroupSim

Specificity Determining Positions (SDPs) are positions in a MSA that are differentially conserved in different functional subgroups^50^. GroupSim calculates the possibility of each position on the input MSA being an SDP. The number of subgroups, and the subgroup each sequence belongs to of the input MSA are user-defined. We performed GroupSim analysis on a combination of FunFams with the same GO annotation (i.e., GMPS or NADS), each FunFam was defined as a subgroup. On each column of the MSA, GroupSim measures the average similarity between pairs of residues within a FunFam, and the average similarities between FunFams. The average within-FunFam similarity subtracted by average between-FunFam similarity is the column score that represents the likelihood of this position to be an SDP^50^. The SDP score threshold of 70 (out of 100) was applied as a significant probability for a position to be an SDP.

### Searching the structural motif templates against databases using ASSAM

The Amino acid pattern Search for Substructures and Motifs (ASSAM) webserver^37^ was recruited to verify the identified structural motifs. The ASSAM algorithm relies on a graph theoretical approach to scan the input motif against structures of protein structure databases^37^, including the CATH database. Databases for HUP structures from the PDB classified by CATH and those predicted by AF2 were generated specifically for this work and have been made available as database options on the Graph theoretical Applications for Substructure Searching (GrAfSS) webserver^58^. ASSAM uses representations of the amino acid side chains as pseudo-atoms to generate vectors, ultimately representing structures as a graph consisting of multiple vectors^37^. The output of ASSAM is a table of all structures matching the input motif, and their RMSD between the input. For the ASSAM outputs we calculated specificity, sensitivity, precision, and accuracy by following equations.

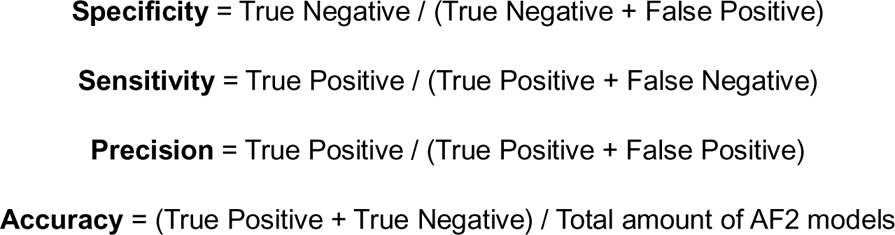

### Phylogenetic tree construction

We used the software MEGA11^71^ to construct the phylogenetic tree based on ATP-PPase domain sequences. A neighbour-joining tree^82^ was built with 1,000 replicates of Bootstrap test^83^. The resulting (bootstrap-consensus) phylogenetic tree was visualized by the software FigTree V1.4.4 (http://tree.bio.ed.ac.uk/software/figtree/).

