## Supplementary Figures for "Understanding structural and functional diversity of ATP-PPases using protein domains and functional families in CATH database"

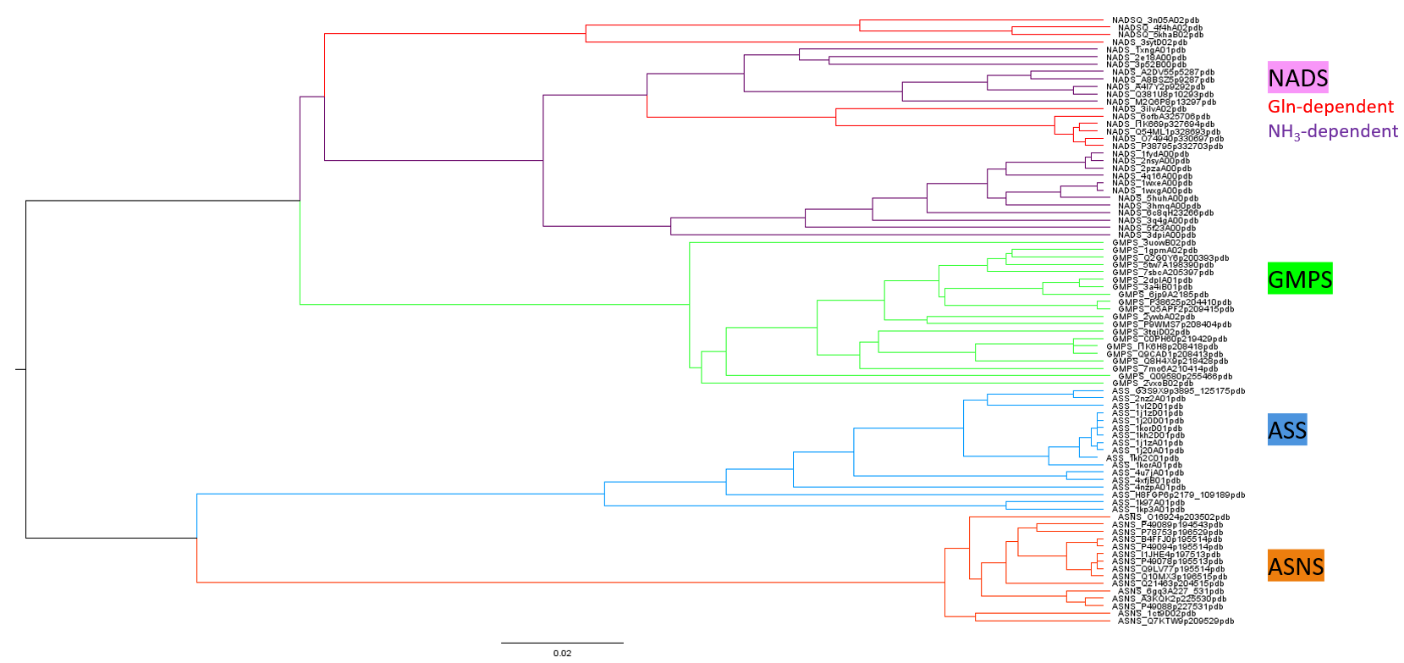

**Figure.S1 UPGMA clustering of ATP-PPase guided by MSTA.** The dendrogram was generated by DendroUPGMA webserver.

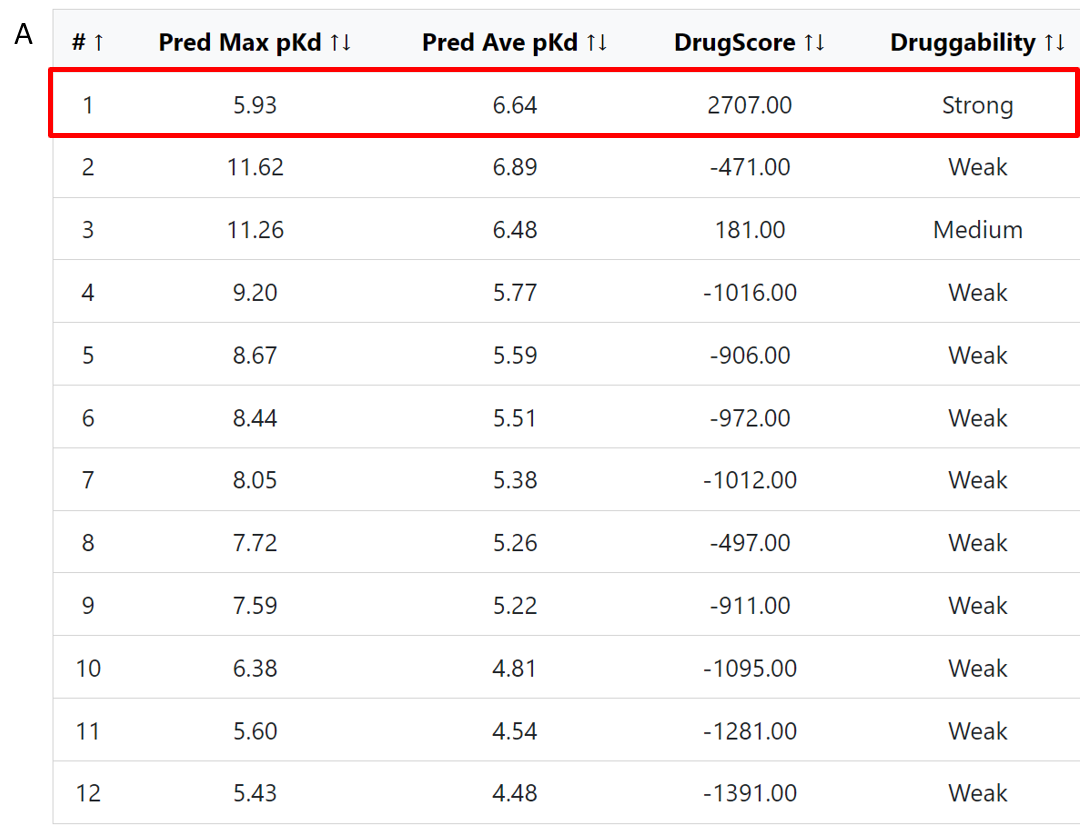

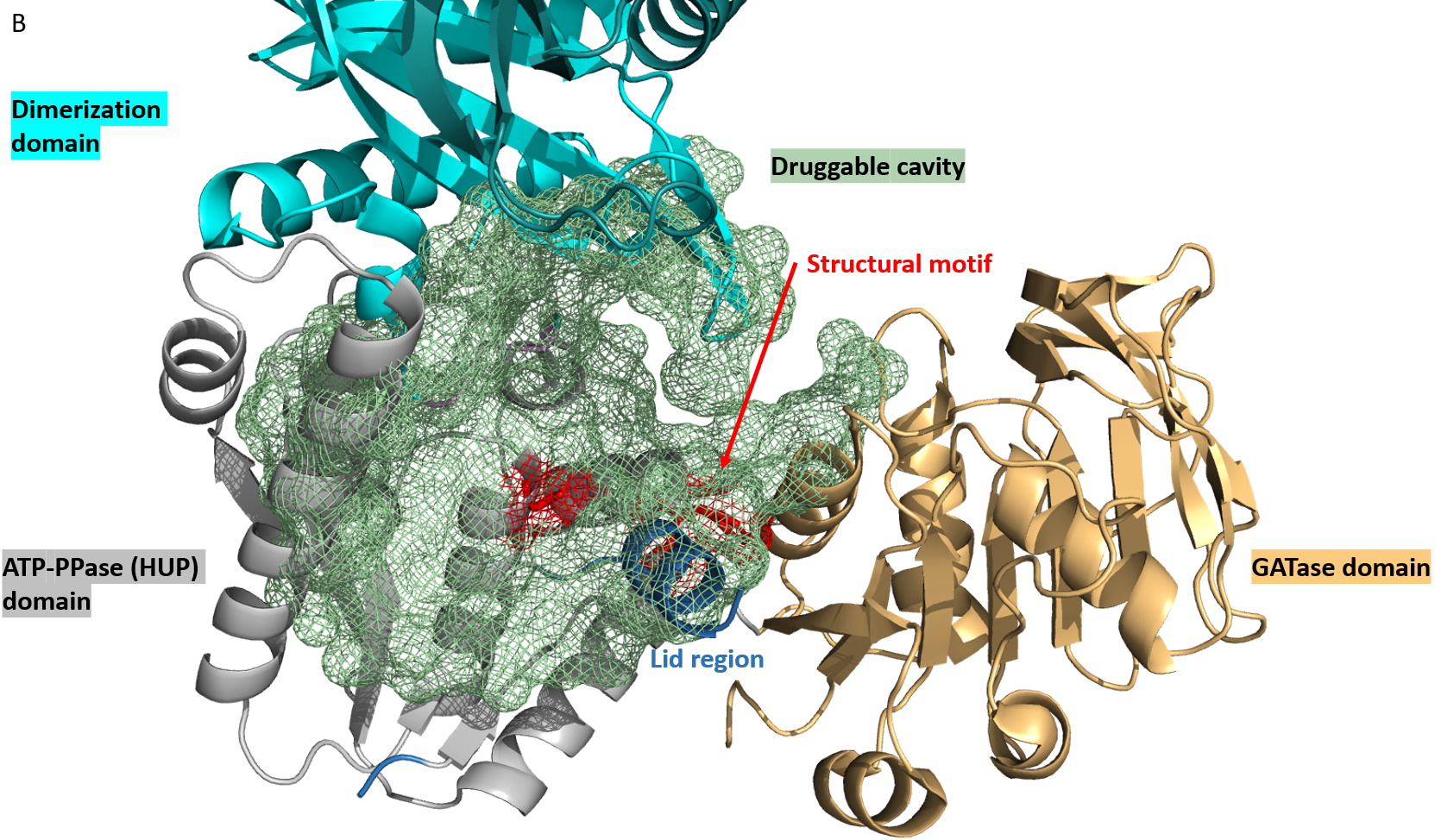

**Figure.S2 CavityPlus predicted cavities of bacterial GMPS.** PDB: 1gpm. (a) Results of all cavities listed on the CavityPlus^1,2^ webserver. The Bacterial GMPS FunFam motif DYLF is found in cavity 1 (red box). (b) Cavity 1 (light green mesh) mapped to the *E.coli* GMPS.

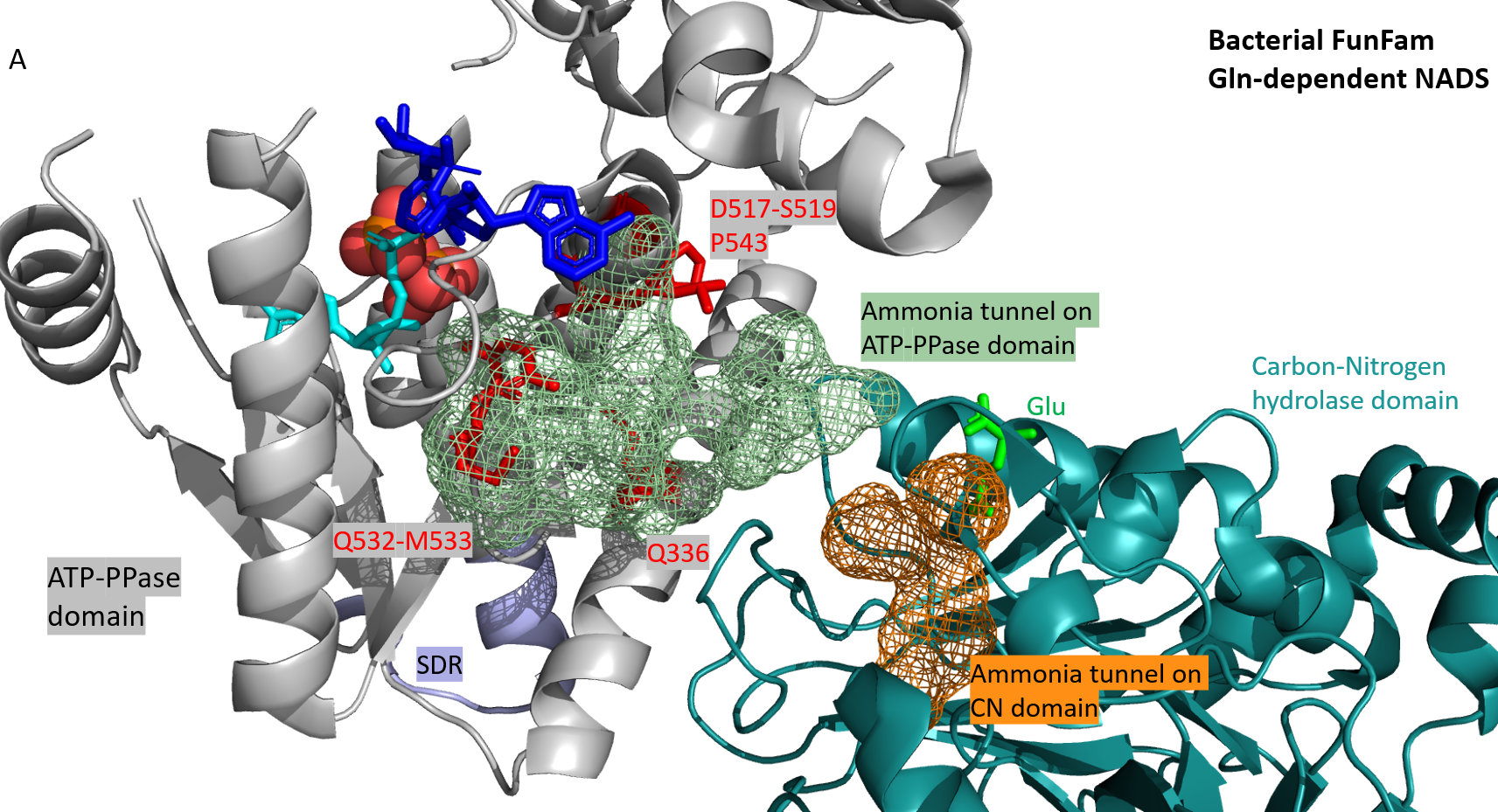

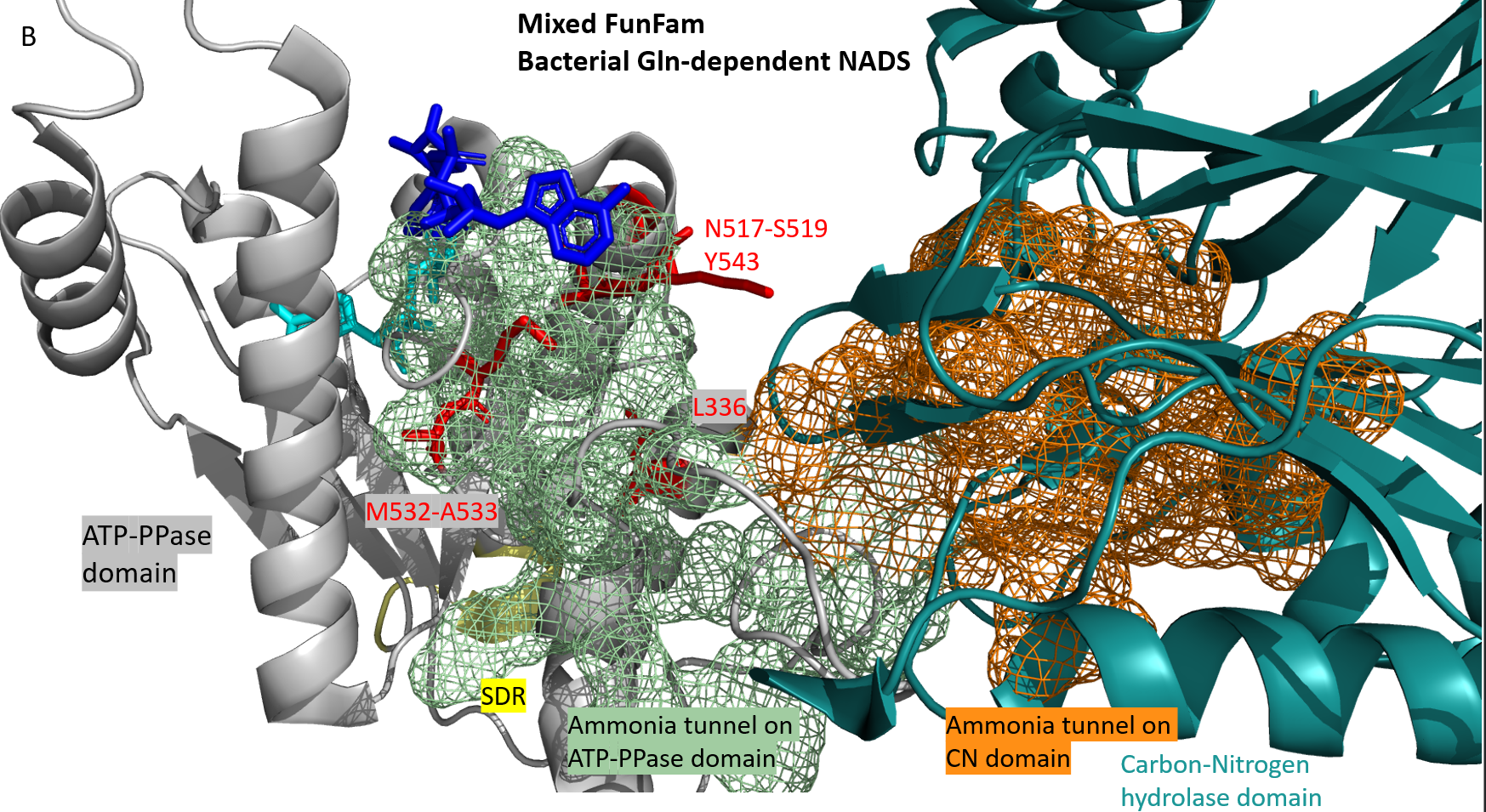

**Figure.S3 Ammonia tunnel and tunnel-interacting SDP residues mapped on bacterial Gln-NADS from two FunFams.** (a) FunFam155 (Bacterial Gln-NADS) member (PDB: 3syt)^3^. (b) FunFam106 (Mixed FunFam) member (PDB: 4f4h)^4^.

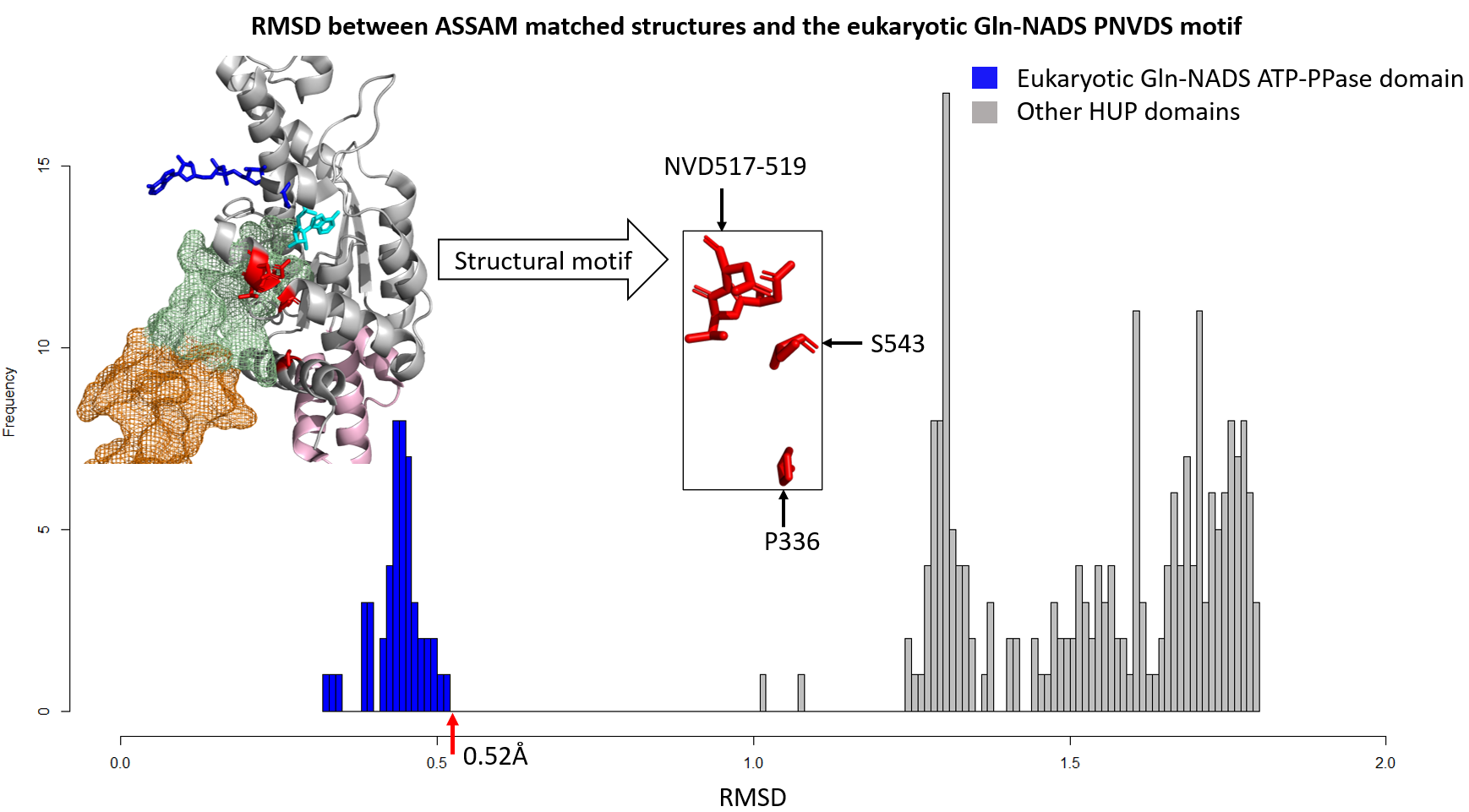

**Figure.S4** **Results from searching the eukaryotic Gln-NADS specific structural motif against a dataset of HUP structures using ASSAM.** PDB: 6ofb. Colour scheme: Light pink-NADS FunFam-specific SDR; Blue-NaAD; Cyan-AMP; Red-SDP residues; Light green and orange mesh-ATP-PPase- and CN-domain portion of NH_3_-tunnel^5^.

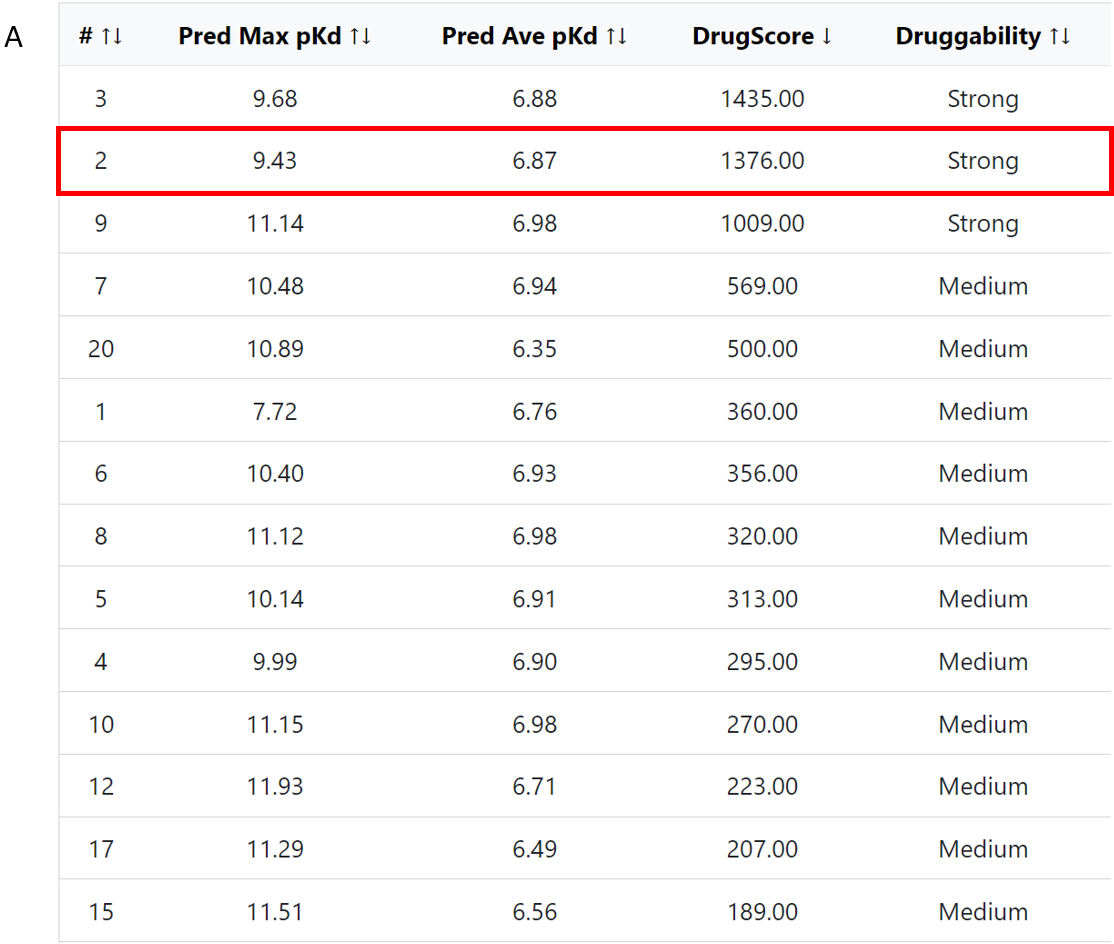

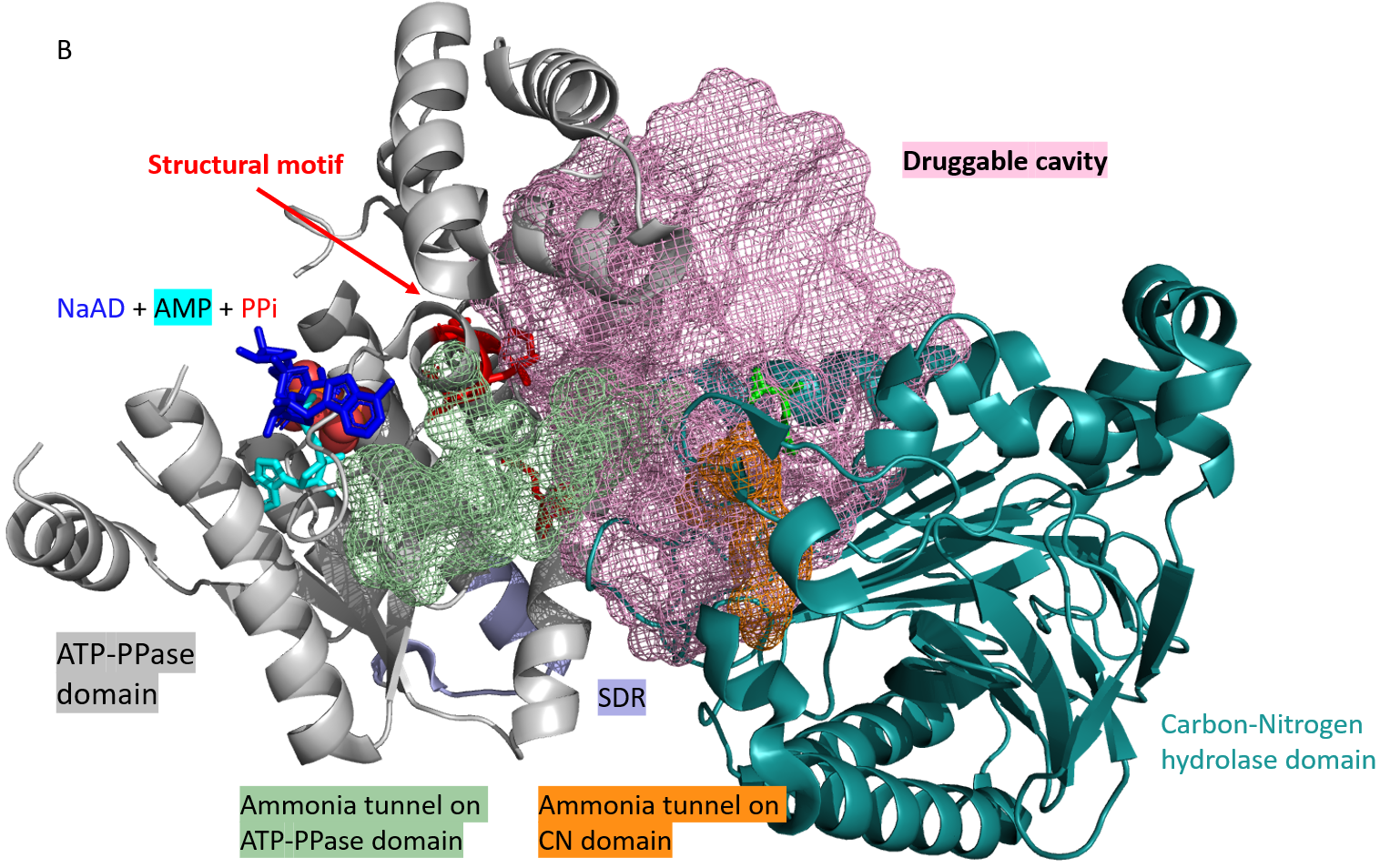

**Figure.S5 CavityPlus predicted cavities of bacterial NADS.** PDB: 3syt. (a) Results of high-druggability cavities listed on the CavityPlus^1,2^ webserver. The Bacterial NADS FunFam motif QDLSP is found in cavity 2 (red box). (b) Cavity 2 (pink mesh) mapped to the *M.tb* NADS.

1.	Xu, Y., Wang, S., Hu, Q., Gao, S., Ma, X., Zhang, W., Shen, Y., Chen, F., Lai, L., and Pei, J. (2018). CavityPlus: a web server for protein cavity detection with pharmacophore modelling, allosteric site identification and covalent ligand binding ability prediction. Nucleic acids research *46*, W374-W379.

2.	Wang, S., Xie, J., Pei, J., and Lai, L. (2023). CavityPlus 2022 Update: An Integrated Platform for Comprehensive Protein Cavity Detection and Property Analyses with User-friendly Tools and Cavity Databases. Journal of Molecular Biology, 168141.

3.	Chuenchor, W., Doukov, T.I., Resto, M., Chang, A., and Gerratana, B. (2012). Regulation of the intersubunit ammonia tunnel in Mycobacterium tuberculosis glutamine-dependent NAD+ synthetase. Biochemical Journal *443*, 417-426.

4.	Santos, A.R.S., Gerhardt, E.C.M., Moure, V.R., Pedrosa, F.O., Souza, E.M., Diamanti, R., Högbom, M., and Huergo, L.F. (2018). Kinetics and structural features of dimeric glutamine-dependent bacterial NAD+ synthetases suggest evolutionary adaptation to available metabolites. Journal of Biological Chemistry *293*, 7397-7407.

5.	Chuenchor, W., Doukov, T.I., Chang, K.-T., Resto, M., Yun, C.-S., and Gerratana, B. (2020). Different ways to transport ammonia in human and Mycobacterium tuberculosis NAD+ synthetases. Nature communications *11*, 1-12.
